# Prolonged loss of D1 receptor signaling promotes maladaptive plasticity of direct pathway striatal projection neurons

**DOI:** 10.64898/2026.11.18.767715

**Authors:** Jenna R. Martin, Saige K. Power, Anne E. Cavanagh, Marta Maltese, Riccardo Melani, Tanya Sippy, Jeremy S. Dasen, Un Jung Kang, Nicolas X. Tritsch

**Affiliations:** Neuroscience Institute, New York University Grossman School of Medicine, New York, NY; Fresco Institute for Parkinson’s and Movement Disorders, New York University Langone Health, New York, NY, U.S.A; Department of Psychiatry, New York University Grossman School of Medicine, New York, NY, U.S.A; Department of Psychiatry, Douglas Hospital Research Centre, McGill University, Montreal, QC, Canada; Integrated Program in Neuroscience, Montreal Neurological Institute, McGill University, Montreal, QC, Canada

## Abstract

As Parkinson’s disease (PD) progresses and nigrostriatal dopamine (DA) neuron degeneration worsens, DA replacement therapies shift from restoring motor function to evoking dyskinesia. How disease progression mediates this transition remains poorly understood. Here, we optimized a 6-hydroxydopamine (6OHDA) model that reliably generates multiple severities of DA axon denervation within the striatum of individual mice. We identify a critical denervation threshold at which direct-pathway striatal projection neurons (dSPNs), which are key drivers of dyskinesia become pathologically hyperresponsive to DA. Using slice electrophysiology, we establish that denervation-induced increase in intrinsic excitability is insufficient to explain the aberrant response of dSPNs to DA. We further demonstrate using genetic and chemogenetic manipulations that prolonged loss of D1 receptor signaling drives the pathological transformation of dSPNs cell-autonomously, and that sustained G_s_-coupled metabotropic receptor signaling in dSPNs is sufficient to both prevent and reverse it. These findings provide a mechanistic framework for understanding how PD progression renders striatal circuits vulnerable to DA-evoked motor complications and identify sustained signaling by G_s_-coupled receptors as a potential avenue for preventing or reversing denervation-induced maladaptive striatal plasticity.

**In Brief (Highlights):**

- dSPN hyperactivation in response to DA treatments emerges only when local DA axons are completely lost
- Increased dSPN intrinsic excitability emerges at intermediate denervation and, as such, cannot explain the shift in DA responsiveness when denervation is complete
- Cell-autonomous deletion of D1 receptors reproduces the pathological shift in dSPN responsiveness to G_s_-coupled metabotropic signaling
- Sustained G_s_-coupled metabotropic signaling prevents and reverses denervation-induced maladaptive plasticity in dSPNs

## Introduction

Parkinson’s disease (PD) is a progressive neurodegenerative disorder characterized by slowing of movements, muscle rigidity and tremor ^1^. These symptoms are caused by the gradual, axon-first degeneration of dopamine (DA)-producing neurons in the substantia nigra pars compacta (SNc) and the subsequent dysregulation of neural activity in dorsal striatal regions that normally receive SNc axons. It is estimated that motor symptoms develop when as many as 60–80% of nigrostriatal axon terminals have degenerated ^2^. Because PD is a progressive disease, striatal denervation continues to worsen post-diagnosis, reaching up to 100% within the following 5 to 10 years ^3^. Striatal activity and motor function can be effectively restored by elevating DA levels with levodopa – the metabolic precursor for DA – or by directly stimulating DA receptors with pharmacological agonists. Later in the disease, however, these same treatments acutely evoke abnormal striatal activity and uncontrolled movements called dyskinesia that severely lower quality of life ^4–6^. This is a significant problem, as the majority of PD patients eventually develop this debilitating motor complication, typically within 5 to 10 years of diagnosis ^4,6^. How DA replacement therapies shift over time from providing therapeutic relief to triggering dyskinesia remains incompletely understood. DA treatments trigger dyskinetic movements within minutes of drug intake by stimulating striatal DA receptors ^7,8^. Although the dose and cumulative history of DA treatments are important factors influencing the expression of dyskinesia (i.e., the severity and duration of each dyskinetic episode) ^4,9–11^, accumulating evidence suggests that the initial emergence of motor complications over the course of the disease (i.e., when DA treatments first evoke dyskinetic movements in the years that follow PD diagnosis) is most strongly linked to the progression and severity of DA loss ^12–15^. In humans, dyskinetic movements first manifests in the limb most affected by the disease, contralateral to the striatum with the greatest loss of DA axons ^4,15–17^. Longitudinal studies show that postponing treatment at the time of PD diagnosis does not delay the emergence of dyskinesia ^18–20^ and, in rare cases of severe and accelerated degeneration following poisoning with the DA neurotoxin MPTP, dyskinesia arises within days of treatment onset ^21–23^. In rodents, levodopa and DA receptor agonists do not evoke dyskinesia if nigrostriatal axons are intact or only partially lesioned – irrespective of treatment dose or duration – but readily do within minutes of first-ever exposure in animals in which SNc lesions are complete ^24–29^. The severity of the loss of nigrostriatal axons is therefore a key determinant of how DA medications affect striatal activity and movement, yet experiments in rodents are rarely designed to compare and contrast multiple levels of denervation. Instead, comparisons are typically made between subjects in which DA axons are either intact or severely lesioned, making it difficult to determine precisely when cellular and molecular phenotypes emerge during the progression of PD.

An important way that SNc degeneration modifies how striatal circuits respond to DA replacement therapies is by disrupting the dynamics of extracellular DA ^30^. In the healthy brain, SNc axons control DA’s extracellular concentration and diffusion by balancing vesicular release with transporter-mediated DA reuptake ^31^. Without the latter, levodopa-derived DA can reach abnormally high levels ^32^. In addition, serotonergic axons substitute for SNc in the parkinsonian striatum to convert levodopa to DA, imposing patterns of release that may not be physiological. However, these ‘presynaptic’ dysregulations are not necessary for the expression of dyskinesia, as DA receptor agonists are equally effective at evoking motor complications ^33–35^.

Another consequence of SNc axon loss is aberrant plasticity of ‘postsynaptic’ circuits. The striatum consists mostly of spiny projection neurons (SPNs) expressing either D1 metabotropic receptors (known as direct-pathway SPNs, or dSPNs) that couple to the G_s_ family of heterotrimeric G proteins (G_olf_ in the case of dSPNs) or D2 metabotropic receptors (known as indirect-pathway SPNs, or iSPNs) that couple the G_i_ family of G proteins, as well as small populations of interneurons that also express DA receptors ^36,37^. Although all modify their electrophysiological properties following SNc lesions and contribute to the expression of dyskinesia in one form or another ^29,38–45^, several lines of evidence point to the importance of maladaptive plasticity in D1 receptor-expressing dSPNs for triggering the profound switch in the brain’s response to DA during the progression of PD ^14^: under normal conditions, dSPNs are sparsely active and resistant to DA-evoked hyperactivation, but following loss of SNc axons, most become aberrantly hyperactive within minutes of systemic administration of levodopa or receptor agonists ^46–51^. iSPNs have not been shown to drastically modify their responsiveness to DA between intact and lesioned brains ^48,49^. Importantly, dSPN hyperactivation is sufficient to evoke dyskinesia, even when SNc axons are intact ^50–55^, and striatal D1 receptors are required for DA medications to evoke dyskinesia in rodents ^56–58^. dSPNs may therefore serve as a common effector through which diverse striatal abnormalities implicated in dyskinesia converge to influence behavior. In line with this, recent studies suggest that the effects of iSPN or D2 receptor manipulations on movement are mediated in part through dSPNs, potentially via recurrent connectivity within the striatum ^49,59–61^. Collectively, these observations suggest that when SNc axon loss reaches a critical threshold, striatal circuits undergo pathological adaptations that fundamentally alter how dSPNs respond to DA replacement, rendering them aberrantly prone to hyperactivation. Understanding precisely when and how this denervation-induced maladaptive plasticity emerges during the progression of PD is therefore critical for developing approaches that prevent motor complications while preserving the therapeutic efficacy of DA replacement.

Toward this end, we established and characterized a 6-hydroxydopamine (6OHDA) model in which multiple severities of SNc axon denervation are reliably generated within the dorsal striatum of individual mice. This approach allowed us to identify the critical degree of denervation needed for dSPNs to profoundly alter their responsiveness to DA *in vivo*: dSPNs became aberrantly hyperactive in response to DA only when all local DA axons were lost. Using slice electrophysiology, we show that the commonly observed increase in dSPN intrinsic excitability emerges at intermediate levels of denervation, indicating that it cannot account for the profound shift in DA responsiveness that occurs with complete denervation. Using genetic and chemogenetic manipulations, we further demonstrate that prolonged loss of D1 receptor signaling in dSPNs drives their pathological transition cell-autonomously, and that sustained signaling by G_s_-coupled metabotropic receptors is sufficient to both prevent and reverse denervation-induced adaptations that render dSPNs hyperresponsive to DA.

## Results

### Intrastriatal 6OHDA infusion produces gradients of DA axon denervation

We first sought an experimental preparation in which dSPNs exposed to varying severities of SNc axon denervation can be quantitatively compared within the same animal. To accomplish this, we slowly infused the DA neurotoxin 6-hydroxydopamine (6OHDA) in the dorsolateral striatum of adult mice to promote axon-first degeneration of SNc neurons^25^ (**Figure 1A**). Because striatal neurons sense DA locally through their dendrites, we quantified DA axon denervation from the perspective of individual neurons. To do so, we immunostained fixed coronal sections for the DA axon marker DAT (**Figure 1B**), obtained high-resolution confocal images tiling the entire dorsal striatum (**Figure 1C, 1D**), seeded 50 µm-diameter region of interests (ROIs) throughout, and for each ROI calculated the fraction of the neuropil innervated by DAT^+^ DA axons (see methods; **Figure 1E**). 6OHDA infusions created gradients of decreasing DA axon density along the mediolateral axis that differed significantly from sham-injected controls in the contralateral hemisphere (*N* = 19 mice; **Figure 1C–1F**). To correct for subtle differences in injection coordinates and volumes between subjects, we plotted DA axon density relative to the denervation boundary, defined as the line separating striatal regions innervated by as little as one DA axon from regions devoid of DA axons entirely (**Figure S1**). We then normalized density values to sham lesions in the contralateral hemisphere, and converted DA axon density to percent denervation (**Figure 1G**). The denervation gradients produced cover a range of severities relevant to the progression of PD, albeit over striatal space, not time. This is advantageous, as it allows us to control for DA axon degeneration as an experimental variable within individual subjects. Because the dendritic fields of dSPNs are small relative to DA gradients ^62^, we hypothesized that this model would allow us to identify denervation severities that cause dSPNs to modify their responsiveness to DA treatments with cellular resolution.

**Figure 1:**
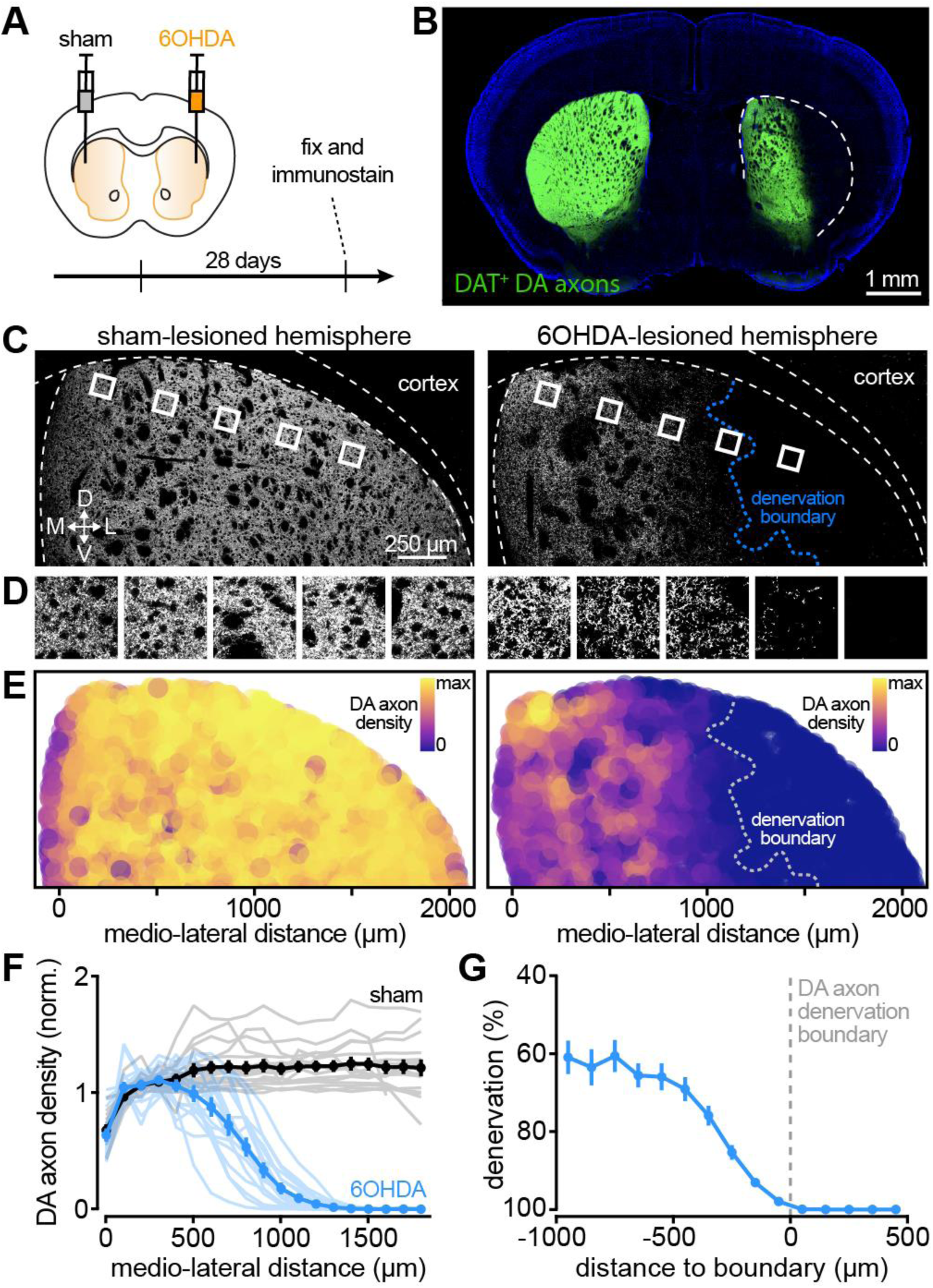
6OHDA produces gradients of DA axon denervation across striatum. A. Diagram of experimental design and timeline. Vehicle (gray) and 6OHDA (orange) are infused in the left and right dorsolateral striatum, respectively. B. Example epifluorescence image of fixed coronal forebrain section immunostained for DAT showing loss of DA axons in the dorsolateral striatum infused with 6OHDA. *Blue*: DAPI nuclear stain. C. Example confocal images of DA axons immunostained for DAT throughout dorsal striatum. *Dotted blue line*: denervation boundary. D. Higher magnification view of insets in (C) at different locations along the medio-lateral axis. E. Color-coded local DA axon density measurements of sham (*left*) and 6OHDA-lesioned (*right*) striatum from (C). F. Mean (± s.e.m.) DA axon density normalized to medial striatum along medio-lateral axis of sham (*black*) and 6OHDA-lesioned (*blue*) striatum (*N =*19 mice). Individual mice shown in gray. G. Mean (± s.e.m.) DA axon denervation severity relative to denervation boundary (*N =* 19 mice).

### dSPNs do not modify their responsiveness to DA until local denervation is severe

We next asked where, along DA axon gradients, dSPNs first undergo denervation-induced maladaptive plasticity rendering them abnormally prone to DA-evoked hyperactivation. To do so, we performed sham and 6OHDA lesions in the left and right dorsolateral striatum of transgenic mice expressing tdTomato in dSPNs. Four weeks later, we tested how dSPNs acutely respond to DA replacement as a function of local DA axon density by administering a single dyskinesiogenic dose of levodopa (5 mg/kg; *N* = 5 mice) or saline (*N* = 7 mice) 90 min before immunostaining striatal sections for DAT and the immediate early gene product c-Fos as a proxy for neuronal activity ^63^ (**Figure 2A**).

**Figure 2:**
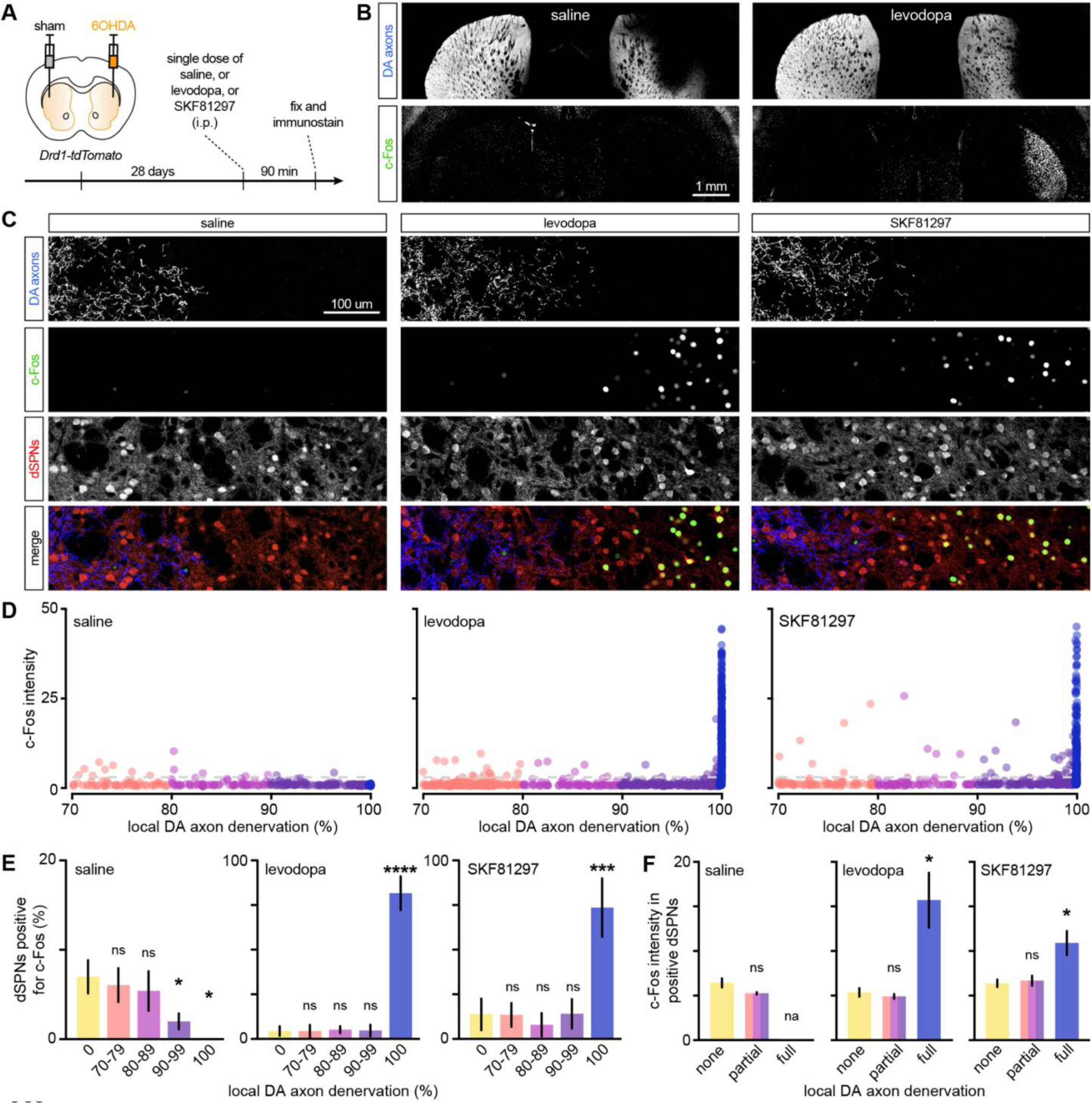
dSPNs do not modify their responsiveness to DA until local denervation is severe. A. Experimental design and timeline. B. Epifluorescence image of dorsal forebrain immunostained for DAT (*top*) and c-Fos (*bottom*) after a single, first-ever challenge with saline (*left*) and levodopa (*right*). Note the exceedingly low expression of c-Fos at baseline and strong induction in the dorsolateral striatum where DA axons are lesioned. C. Confocal images of DAT, c-Fos and tdTomato fluorescence (individual channels shown as gray scale, merged in color) straddling the DA axon denervation boundary in the 6OHDA-lesioned striatum of *Drd1-tdTomato* mice challenged with saline (*left*), levodopa (5 mg/kg; *middle*) or SKF81297 (3 mg/kg; *right*). Note the strong induction of c-Fos in dSPNs in regions devoid of DA axons. D. Intensity of nuclear c-Fos fluorescence (normalized to background neuropil) in dSPNs exposed to different local DA axon denervation in the dorsal striatum of example mice challenged with saline (*left*), levodopa (*middle*) or SKF81297 (*right*). Colors denote DA axon denervation bin (70-79%, 80-89%, 90-99% and 100%). E. Mean (± s.e.m.) percentage of all dSPNs considered positive for c-Fos at different local DA axon denervation across all subjects following challenge with saline (*N =* 7; one-way ANOVA, *p* = 0.0072; Dunnett’s *post-hoc* tests: 0 vs. 90-99, *p* = 0.0451; 0 vs. 100, *p* = 0.0283), levodopa (*N =* 5; *p* < 0.0001; 0 vs. 100, *p* < 0.0001) or SKF81297 (*N =* 7; *p* < 0.0001; 0 vs. 100, *p* = 0.0001). F. Same as (E) for mean intensity of nuclear c-Fos fluorescence in dSPNs deemed positive for c-Fos in mice challenged with saline (*p* = 0.13, Wilcoxon signed rank test), levodopa (*p* = 0.035, mixed-effects model; Dunnett’s *post-hoc* test: none vs. full, *p* = 0.014) and SKF81297 (*p* = 0.027, Mixed-effects model; Dunnett’s *post-hoc* test: none vs. full, *p* = 0.029). DA axon denervation binned as none (0%), partial (70-99%) and full (100%). Too few dSPNs in fully-lesioned striatum were c-Fos^+^ in saline-treated mice for quantification.

In saline-treated mice, only a small fraction of tdTomato^+^ dSPNs (5–10 %; *n* = 1,279 dSPNs) were considered positive for c-Fos, consistent with their sparse activation and low firing rates (**Figure 2B**). As the degeneration of DA axons increased in severity along 6OHDA gradients, the number of dSPNs positive for c-Fos decreased further, confirming their hypoactivation in PD (**Figures 2C– E**).

In mice challenged with levodopa, we observed a marked increase in both the number of c-Fos^+^ cells and the intensity of c-Fos immunolabeling selectively in the dorsolateral striatum of the lesioned hemisphere, where DA axon denervation is most severe (**Figure 2B**). The vast majority of these cells (96.6 ± 0.4 %) were tdTomato^+^ dSPNs (**Figures 2C, S2A**), consistent with aberrant dSPN hyperactivation in response to a first-ever levodopa challenge^35,46–49^. We observed similar levodopa-evoked c-Fos induction in DA-denervated regions of mice injected with 6OHDA into the dorsolateral striatum (*N* = 3 mice; **Figure S2B**) or the medial forebrain bundle (*N* = 2 mice; **Figure S2C**), indicating that the c-Fos response is neither specific to the dorsolateral striatum, nor attributable to local effects of 6OHDA infusion, but rather tracks DA axon loss.

Quantification of high-resolution confocal images revealed that the percentage of dSPNs expressing c-Fos in levodopa-challenged mice only increased in striatal regions without detectable DA axons, reaching 81 ± 4 % of dSPNs (**Figures 2C–E**). The immunolabeling intensity of c-Fos^+^ dSPNs also increased significantly in denervated regions (**Figure 2F**). Strikingly, c-Fos induction closely followed the contours of the denervation boundary, reaching a plateau within 150 μm of the nearest visible DA axon (**Figures 2C, S2D–S2F**). By contrast, levodopa restored c-Fos expression to near-control levels in dSPNs on the other side of the denervation boundary, wherever some DA axon innervation was maintained (90-99% denervation; **Figure 2E**). These findings indicate that the manner in which dSPNs respond to levodopa is strongly dependent on the local density of DA axons within striatum.

Because c-Fos expression does not necessarily reflect neuronal activity ^64,65^, we re-analyzed a functional Ca^2+^ imaging dataset we previously published, in which the activity of dSPNs was monitored longitudinally *in vivo* before and after 6OHDA lesioning of the SNc ^48^ (**Figures S2G– S2J**). We stratified fields of view according to whether *post-hoc* analyses detected residual DA axons in dorsal striatum or not (**Figure S2I**). Although dSPNs in both groups became comparably hypoactive following SNc degeneration, their response to acute levodopa differed markedly: the number of active dSPNs returned to pre-lesion levels in fields of view with a few DA axons remaining, but increased beyond the range observed under normal conditions in regions devoid of DA axons (**Figure S2J**). These data provide functional evidence that local DA axon innervation predicts whether levodopa restores dSPN activity toward normal levels or instead evokes pathological hyperactivity.

Denervation severity could alter the effects of levodopa presynaptically, by modifying the levels and dynamics of levodopa-derived DA, or postsynaptically, by inducing maladaptive plasticity that alters the responsiveness of striatal circuits to DA. To distinguish between these possibilities, we bypassed the requirement for levodopa conversion to DA by directly stimulating D1 receptors using the selective agonist SKF81297 (3 mg/kg) in a new cohort of mice (*N* = 7). SKF81297 altered c-Fos expression in dSPNs in a pattern similar to levodopa: it restored c-Fos expression toward control levels in regions with severe but incomplete denervation, while inducing pathological c-Fos expression in regions with no detectable DA axons, closely following the denervation boundary (**Figures 2C–2F**). Similar results were also obtained by staining for Arc (**Figure S3**), an immediate early gene product more tightly linked to synaptically driven neuronal activation ^66,67^. Together, these findings suggest that DA denervation induces postsynaptic adaptations that fundamentally alter how dSPN respond to DA: dSPNs remain resistant to D1 receptor-mediated hyperactivation for as long as residual DA axon innervation persists, but become highly susceptible to it when local denervation exceeds a critically severe threshold.

### Changes in intrinsic excitability are not sufficient for making dSPNs hyperresponsive to DA

Previous studies show that DA depletion increases the intrinsic excitability of dSPNs ^68–71^. This denervation-induced plasticity might be responsible for making dSPNs aberrantly hyperresponsive to DA when denervation is complete. If so, changes in intrinsic excitability ought to map onto striatal regions lacking DA innervation. To test this, we generated 6OHDA gradients in the striatum of *Drd1-tdTomato* mice (*N* = 20) and measured 4 weeks later the intrinsic excitability of tdTomato^+^ dSPNs exposed to different degrees of DA axon loss using whole-cell current-clamp recordings. Denervation severity was quantified *post-hoc* from confocal images of biocytin-filled neurons using DAT immunofluorescence (**Figure 3A**). In line with previous work ^68–71^, dSPNs in DA-depleted striatal regions (*n* = 53 dSPNs) had similar resting membrane potentials but higher membrane resistances and were significantly more excitable than dSPNs in DA-intact regions (*n* = 44; **Figures 3B–3F**). However, these modifications were already evident at intermediate denervation levels (70-95%; *n* = 42 dSPNs; **Figures 3B–3F**), indicating they are not specific to regions where dSPNs are prone to hyperactivation and, as such, are not sufficient to explain the differential behavior of dSPNs across denervation boundaries.

**Figure 3:**
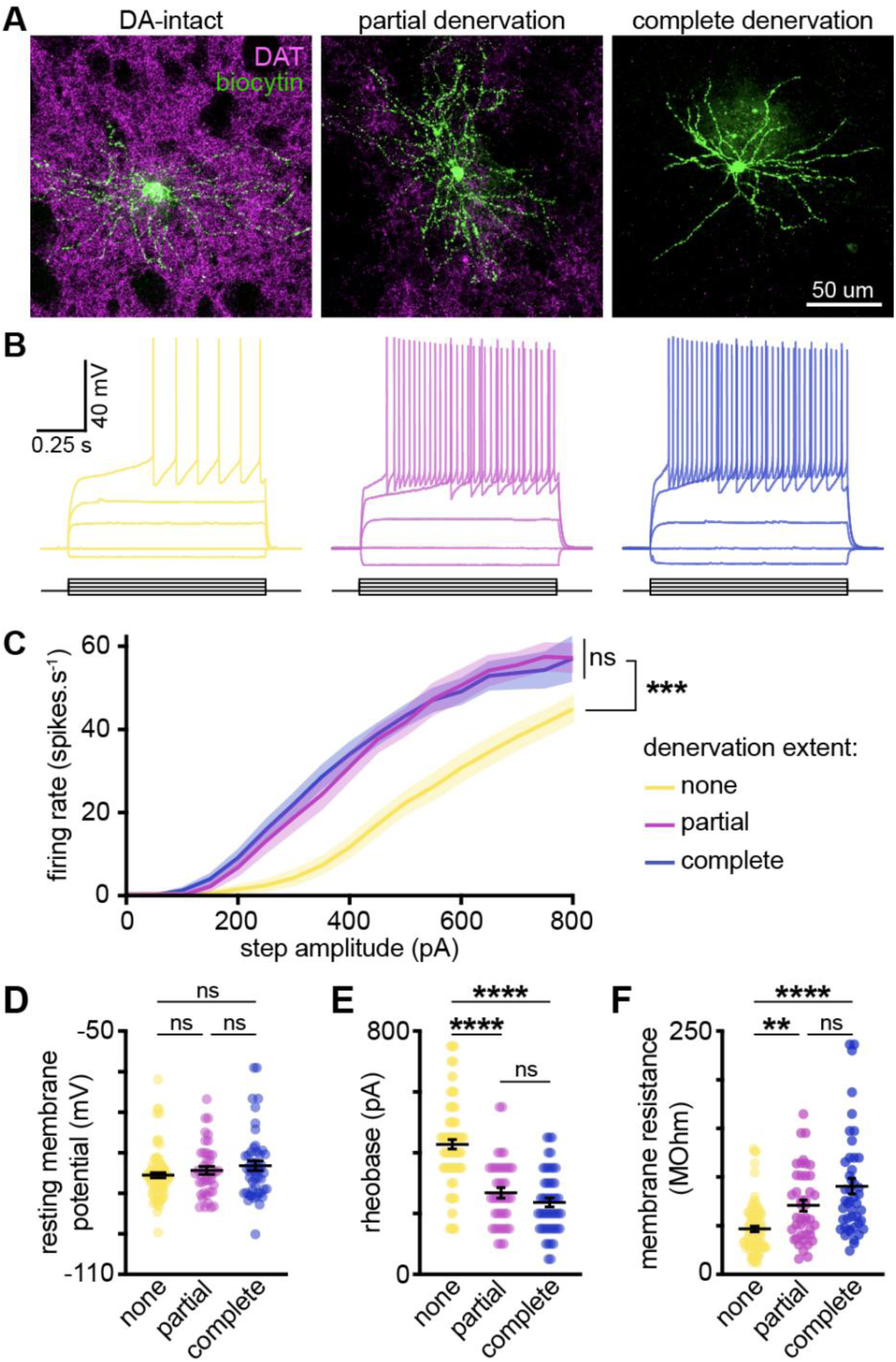
Changes in intrinsic excitability are not sufficient for making dSPNs hyperresponsive to DA. A. Example confocal images of biocytin-filled dSPN (*green*) in in striatal slices from *Drd1-tdTomato* mice with locally intact (*left*), partially denervated (*middle*) and completely denervated (*right*) DAT^+^ DA axons (*purple*). B. Representative voltage responses of dSPNs in DA-intact (*yellow*), partially denervated (*purple*) and completely denervated (*blue*) striatum in response to injection of identical current steps (*bottom*). Note increase in intrinsic excitability in partially and completely denervated dSPNs. C. Mean current–response curves (shaded regions illustrate s.e.m.) for dSPNs locally exposed to different denervation levels (none, *n* = 44 dSPNs; partial (70-95%), *n* = 42 dSPNs; complete, *n* = 53 dSPNs; mixed-effects model for lesion extent: *p* < 0.0001; Tukey’s post-hoc tests: partial vs. complete, *p* > 0.2 for all current steps; partial or complete vs. none, *p* < 0.001 for steps 300-700 pA). D. Resting membrane potential of same dSPNs in C (colored dots: individual cells; mean ± s.e.m. shown in black; *p* = 0.2, Kruskal-Wallis). E. Same as (D) for minimum current required to evoke action potentials (rheobase; *p* < 0.0001, Kruskal-Wallis; Dunn’s *post-hoc* tests: none vs. partial or complete, *p* < 0.0001). F. Same as (D) for dSPN membrane resistance (*p* < 0.0001, Kruskal-Wallis; Dunn’s *post-hoc* tests: none vs. partial, *p* = 0.0011; none vs. complete, *p* < 0.0001).

### Cell-autonomous loss of D1 receptors renders dSPNs aberrantly responsive to G_s_-coupled metabotropic signaling

How do dSPNs sense denervation severity? They may do so cell-autonomously, by detecting declining signals from surface receptors for DA, co-released transmitters or cell-surface molecules on SNc axons ^72–75^, or indirectly in response to lesion-induced circuit-wide dysfunctions ^39,76,77^. Given the striking spatial correspondence between DA denervation boundaries and dSPN hyperactivation, we hypothesized that prolonged loss of D1 receptor signaling in dSPNs cell-autonomously triggers maladaptive plasticity that fundamentally alters how dSPNs respond to the activation of G_s_-coupled metabotropic receptors. If so, selectively removing D1 receptors from sparse populations of dSPNs in the adult brain should be sufficient to render these cells aberrantly responsive to G_s_-coupled metabotropic receptor stimulation, even when local DA axon innervation and the surrounding circuitry remain intact.

To test this, we conditionally knocked out (cKO) D1 receptors from dSPNs by virally expressing Cre in *Drd1*^flox/flox^ mice ^78^. To limit Cre expression to dSPNs and minimize circuit-wide non-cell-autonomous effects, we used a retrograde viral approach that sparsely and selectively targets dSPNs from the substantia nigra pars reticulata (SNr; **Figure 4A**). Importantly, because deleting D1 receptors precludes the use of levodopa or SKF81297 to reveal denervation-evoked maladaptive plasticity in dSPNs, we also transduced Cre-positive dSPNs with the G_s_-coupled chemogenetic actuator rM3Ds ^79^, which previous studies show can substitute for D1 receptors to evoke dSPN hyperactivation and dyskinesia in DA-depleted mice ^52^. To facilitate the identification of Cre-positive dSPNs transduced with rM3Ds in fixed tissue, we generated viral vectors that co-express FLAG-tagged rM3Ds and a HA-tagged nuclear marker (histone 2B, or H2B), both in a Cre-dependent fashion (**Figures S4A–S4B**). Control experiments confirmed our ability to genetically ablate D1 receptors from the striatum of adult mice within 4 weeks of Cre expression (**Figure S4C**), to retrogradely transduce dSPNs from SNr (**Figure S4D**), and to trigger aberrant hyperactivation of rM3Ds-expressing dSPNs in DA-depleted mice following acute administration of the potent chemogenetic activator Deschloroclozapine ^80^ (DCZ; **Figure S4E**).

**Figure 4:**
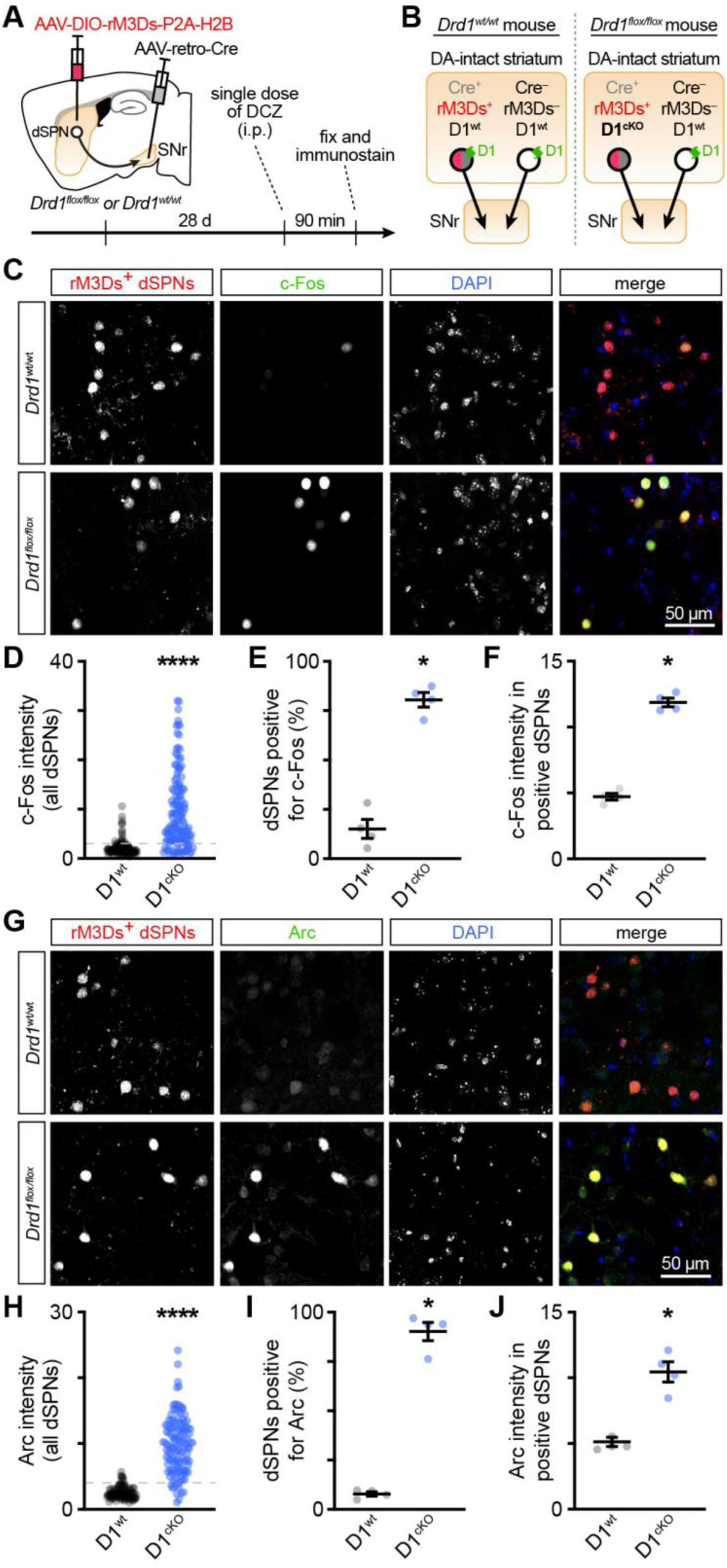
Cell-autonomous loss of D1 receptors renders dSPNs aberrantly responsive to G_s_-coupled metabotropic signaling. A. Experimental design and timeline. Because dSPNs are the only striatal neuronal population projecting directly to the substantia nigra pars reticulata (SNr), they can be selectively targeted for retrograde viral transduction by injecting viral vectors into the SNr (see Figure S4D for validation). B. Schematic of possible experimental conditions obtained from retrograde delivery of Cre to dSPNs via SNr and intrastriatal viral expression of Cre-dependent rM3Ds in *Drd1^wt/wt^*(*left*) and *Drd1*^flox/flox^ mice (*right*). Yellow boxes depict striatum and SNr. Circles and arrows represent dSPNs and their axons. Red and gray fills in dSPNs represent rM3Ds and Cre expression, respectively. D1 receptor expression is shown in green. C. Example confocal images of Cre^+^ dSPNs expressing rM3Ds (*red*; nuclear label for HA-tagged H2B) in the DA-intact striatum of *Drd1*^wt/wt^ (*top*) and *Drd1*^flox/flox^ mice (*bottom*) immunolabeled for c-Fos (*green*). DAPI nuclear stain shown in blue. Note how conditional deletion of D1 receptors in dSPNs leaves dSPNs aberrantly hyperresponsive to rM3Ds stimulation, as revealed by strong c-Fos induction. D. Distribution of c-Fos fluorescence intensity (normalized to background neuropil) in rM3Ds^+^ dSPNs expressing D1 receptors (D1^wt^; *black*) or not (D1^cKO^; *blue*). Dashed line depicts threshold for deeming dSPNs positive for c-Fos (*n* = 33 randomly sampled dSPNs per mouse from *N* = 4 mice per group; *p* < 0.0001, nested t test). E. Mean (± s.e.m.) percentage of D1^wt^ and D1^cKO^ dSPNs considered positive for c-Fos (*N =* 4 *Drd1*^wt/w*t*^ and *Drd1*^flox/flox^ mice each; *p* = 0.0286, Mann–Whitney U test). F. Same as (E) for mean intensity of nuclear c-Fos fluorescence in D1^wt^ and D1^cKO^ dSPNs deemed positive for c-Fos (*p* = 0.0286). G. Same as (C) for Arc immunofluorescence. H. Same as (D) for Arc fluorescence intensity (*n* = 30 randomly sampled dSPNs per mouse from *N =* 4 mice per group; *p* = 0.0005, nested t test). I. Same as (E) for dSPNs positive for Arc (*p* = 0.0286). J. Same as (F) for Arc fluorescence intensity in Arc^+^ dSPNs (*p* = 0.0286).

We therefore applied our experimental protocol to wild-type and *Drd1*^flox/flox^ mice (*N* = 4 each) and, after acute treatment with DCZ, quantified the immunofluorescence intensity of c-Fos and Arc in rM3Ds^+^ dSPNs, which necessarily co-express Cre and thus either possess D1 receptors (D1^wt^) or not (D1^cKO^) depending on genotype (**Figures 4A–4B**). The proportion of rM3Ds^+^ dSPNs that were positive for c-Fos and Arc in DCZ-treated wild-type mice (**Figures 4C, 4E, 4G, 4I**) was similar to that of dSPNs treated with SKF81297 in DA-intact *Drd1-tdTomato* mice (**Figures 2E, S3C**), suggesting that co-expression of Cre, rM3Ds and H2B do not impair dSPN activity. In *Drd1*^flox/flox^ mice, the same DCZ treatment strongly induced c-Fos and Arc expression, such that 83 ± 3 % and 90 ± 5 % of rM3Ds^+^ dSPNs stained positive for c-Fos (**Figure 4C–4F**) and Arc (**Figure 4G–4J**), respectively. This marked increase is similar to that observed in DA-depleted regions of *Drd1-tdTomato* mice following acute treatment with SKF81297 (**Figures 2E–2F, S3C–S3D**), despite DA axons remaining intact in *Drd1*^flox/flox^ mice. These results show that ablating D1 receptors from dSPNs is sufficient to replicate the pathological effects of complete DA axon denervation, and thus, strongly suggest that dSPNs undergo maladaptive plasticity cell-autonomously during the progression of PD when D1 receptor-mediated signaling falls below a critical level.

### Sustained D1-like signaling protects dSPNs from becoming aberrantly hyperresponsive to DA

If loss of D1 receptor-mediated signaling drives the maladaptive shift in dSPN responsiveness to DA, we reasoned that artificially sustaining D1-like signaling might protect dSPNs from this plasticity despite DA axon degeneration. To test this hypothesis, we used a chemogenetic approach, selectively expressing rM3Ds in dSPNs of *Drd1-Cre* mice and administering DCZ via the drinking water to sustain G_s_-coupled metabotropic signaling in dSPNs both before and throughout 6OHDA-mediated denervation of the dorsolateral striatum (**Figures 5A–5B**). In 6OHDA-lesioned mice exposed to regular water (*N* = 5), acute SKF81297 treatment strongly induced c-Fos (**Figure 5C–5E**) and Arc (**Figures 5H–5I**) in 86 ± 8 % and 83 ± 4 % of rM3Ds^+^ dSPNs, respectively, as before (**Figures 2E–2F, S3C–S3D**). By contrast, the SKF81297-mediated induction of c-Fos and Arc in rM3Ds^+^ dSPNs was significantly blunted in mice exposed to DCZ (*N* = 8 for c-Fos and *N* = 6 for Arc; **Figures 5C–5J**), approaching expression levels normally observed in DA-intact mice (**Figure 2E–2F, S3C–S3D**). This effect was specifically mediated by DCZ acting through rM3Ds, as chronic DCZ administration in the drinking water did not significantly affect c-Fos and Arc induction in control mice not expressing rM3Ds in dSPNs (**Figure S4F**). These data demonstrate that sustaining G_s_-coupled metabotropic signaling in dSPNs is sufficient to prevent these cells from becoming aberrantly hyper-responsive to DA when SNc axons are lost.

**Figure 5:**
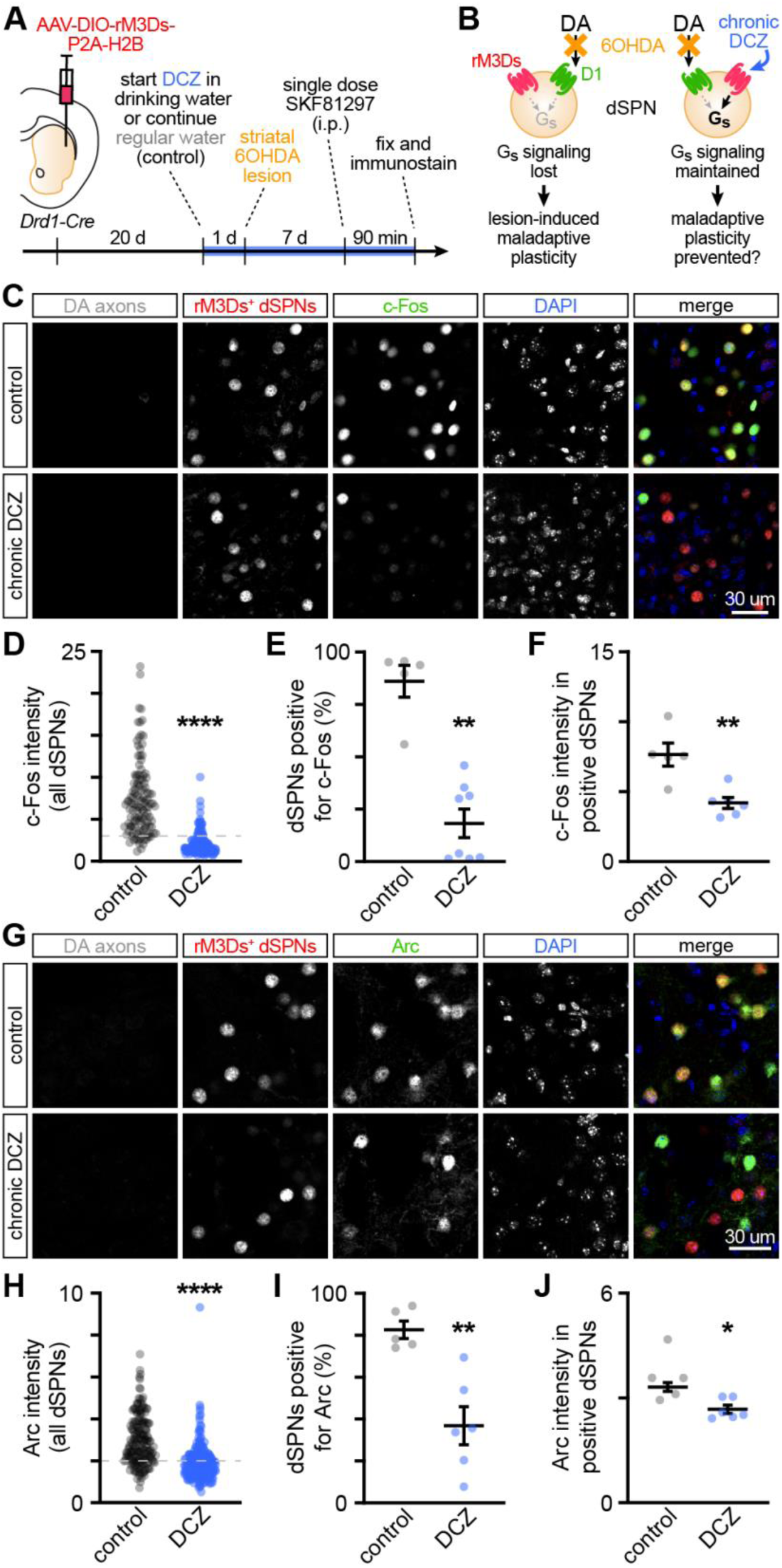
Sustained D1-like signaling protects dSPNs from becoming hyperresponsive to DA. A. Experimental design and timeline. The G_s_-coupled chemogenetic actuator rM3Ds was virally expressed in dSPNs of *Drd1-Cre* mice 3 weeks prior to lesioning DA axons in the dorsolateral striatum. Chemogenetic activation of rM3Ds^+^ dSPNs was initiated 24h prior to lesioning DA axons by supplementing drinking water with DCZ (10 mg/L) to ensure continuous G_s_-coupled metabotropic signaling in dSPNs post-lesion. 7 days is sufficient to reveal lesion-evoked striatal maladaptive plasticity ^35^. B. Schematic of dSPN outlining experimental conditions and hypotheses. *Left*, mice on regular water serve as positive control, confirming that 7 days is sufficient to reveal 6OHDA lesion-induced maladaptive plasticity in dSPNs following a single challenge with SKF81297. *Right*, experimental group provided DCZ in drinking water to test the hypothesis that sustained activation of rM3Ds can substitute for D1 receptor signaling in dSPNs locally deprived of DA. C. Example confocal images of rM3Ds^+^ dSPNs (nuclear label for HA-tagged H2B) in DA-depleted striatum of *Drd1-Cre mice* given DCZ (*bottom*) or not (*top*) in drinking water. DAT immunolabel confirms complete local denervation. Merged image shows rM3Ds^+^ dSPNs in red, c-Fos in green and DAPI in blue. Note strong induction of c-Fos in dSPNs SKF81297 in positive control group (top) and protection of rM3Ds^+^ dSPNs in DCZ-treated mice. D. Distribution of c-Fos fluorescence intensity (normalized to background neuropil) in rM3Ds^+^ dSPNs from mice chronically receiving DCZ (*blue*) or not (control; *black*) via drinking water. Dashed line depicts threshold for deeming dSPNs positive for c-Fos (*n* = 24 randomly sampled dSPNs per mouse from *N =* 5 control mice and *N =* 8 DCZ-treated mice; *p* < 0.0001, nested t test). E. Mean (± s.e.m.) percentage of dSPNs considered positive for c-Fos in control (*N =* 5) and DCZ-treated (*N =* 8) mice (*p* = 0.0016, Mann–Whitney U test). F. Same as (E) for mean intensity of nuclear c-Fos fluorescence in dSPNs deemed positive for c-Fos (*p* = 0.0087). G. Same as (C) for Arc immunofluorescence. H. Same as (D) for Arc fluorescence intensity (*n* = 36 randomly sampled dSPNs per mouse from *N =* 5 control and *N =* 6 DCZ-treated mice; *p* = 0.0030, nested t test). I. Same as (E) for dSPNs positive for Arc (*p* = 0.0043). J. Same as (F) for Arc fluorescence intensity in Arc^+^ dSPNs (*p* = 0.0173).

### Sustained D1-like signaling reverses lesion-evoked dSPN maladaptive plasticity

Having found that sustained rM3Ds activation prevents denervation-induced maladaptive plasticity in dSPNs, we next asked whether sustained G_s_-coupled metabotropic signaling could also reverse this plasticity once established (**Figure 6B**). To test this, we used the same experimental paradigm but delayed DCZ administration until 3 weeks after 6OHDA lesioning of DA axons in the dorsolateral striatum (**Figure 6A**), by which time dSPNs had already undergone the pathological shift in their responsiveness to DA (see **Figure 5**).

**Figure 6:**
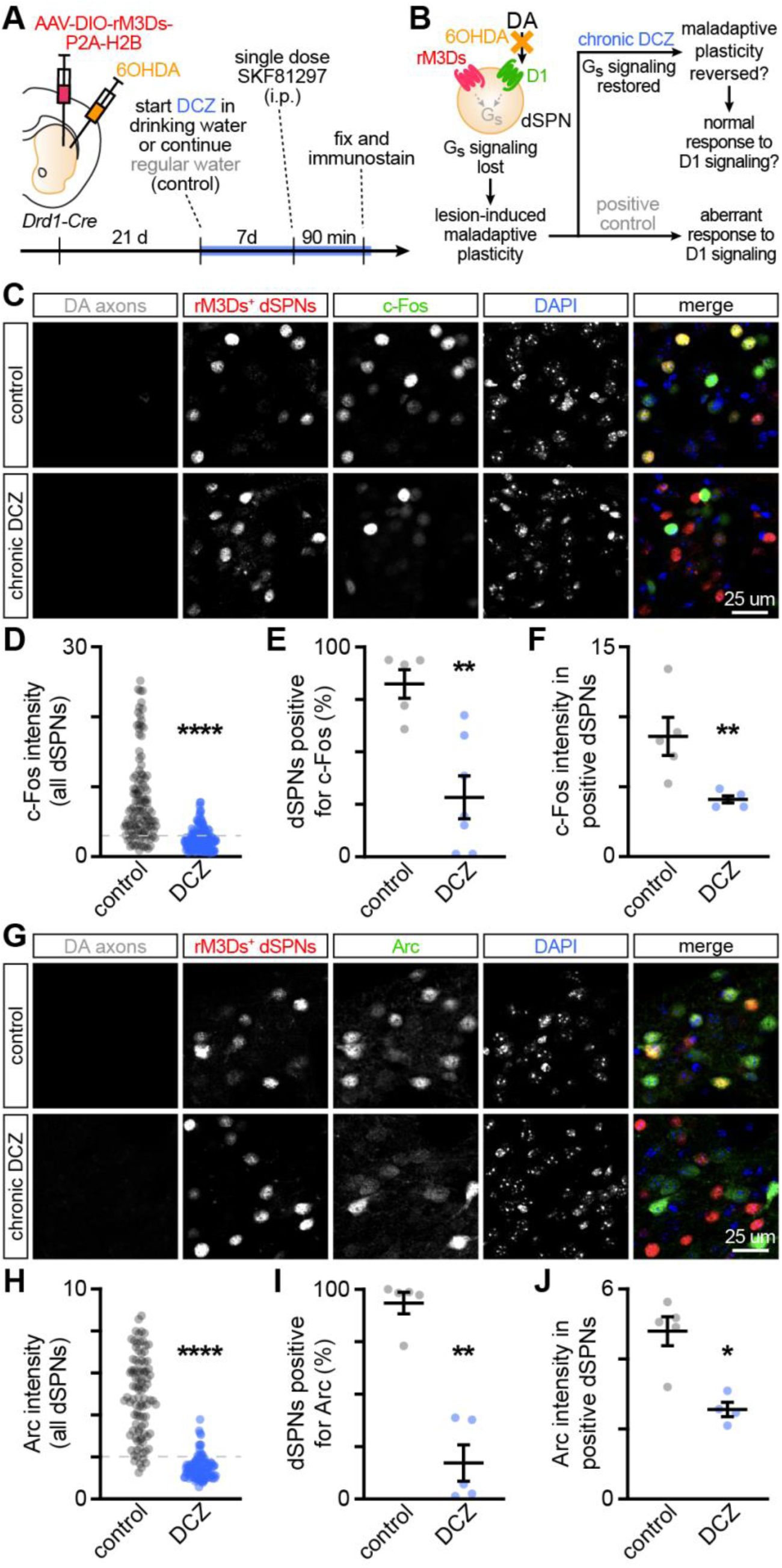
Sustained D1-like signaling reverses lesion-evoked dSPN maladaptive plasticity. A. Design and timeline of experiment testing if sustained chemogenetic activation of rM3Ds in dSPNs can reverse established lesion-induced maladaptive plasticity. B. Schematic of experimental conditions and hypotheses. In this variant of Figure 5, DCZ is added to the drinking water 3 weeks *after* lesioning striatal DA axons with 6OHDA, at which point maladaptive plasticity in dSPNs is already established to test whether restoring G_s_-coupled metabotropic signaling in dSPNs can cell-autonomously reverse maladaptive plasticity and prevent dSPNs from responding aberrantly to DA replacement, even when DA axons are locally lost. Mice on regular water serve as positive control. C. Confocal images of DAT^+^ DA axons (*gray*), rM3Ds^+^ dSPNs (*red*), c-Fos (*green*) and DAPI (*blue*) in a striatal region devoid of DA axons. Under control conditions in mice given regular water (*top*), SKF81297 challenge triggers strong c-Fos induction in dSPNs, confirming maladaptive plasticity secondary to DA axon loss. In mice treated with DCZ (*bottom*), rM3Ds^+^ dSPNs are no longer prone to SKF81297-evoked c-Fos induction. D. Distribution of c-Fos fluorescence intensity (normalized to background neuropil) in rM3Ds^+^ dSPNs from control (*black*) or DCZ-treated (*blue*) mice. Dashed line depicts threshold for deeming dSPNs positive for c-Fos (*n* = 24 randomly sampled dSPNs per mouse from *N =* 5 control and 7 DCZ-treated mice; *p* = 0.0026, nested t test). E. Mean (± s.e.m.) percentage of dSPNs considered positive for c-Fos in control and DCZ-treated mice (*p* = 0.0051, Mann–Whitney U test). F. Same as (E) for mean intensity of nuclear c-Fos fluorescence in dSPNs deemed positive for c-Fos (*p* = 0.0079). G. Same as (C) for Arc immunofluorescence. H. Same as (D) for Arc fluorescence intensity (*n* = 19 randomly sampled dSPNs per mouse from *N =* 5 control and DCZ-treated mice each; *p* = 0.0002, nested t test). I. Same as (E) for dSPNs positive for Arc (*p* = 0.0079). J. Same as (F) for Arc fluorescence intensity in Arc^+^ dSPNs (*p* = 0.0159).

After one week of oral treatment, mice received a single dose of SKF81297 to assess denervation-induced maladaptive plasticity in dSPNs. As expected, SKF81297 robustly increased c-Fos (**Figure 6C–6F**) and Arc (**Figure 6G–6J**) expression in the majority of rM3Ds⁺ dSPNs within fully denervated striatal regions of control mice kept on regular drinking water (*N* = 5). By contrast, in mice treated with DCZ, the expression of c-Fos and Arc in rM3Ds^+^ dSPNs returned to pre-lesion levels (*N* = 7 for c-Fos and *N* = 5 for Arc; **Figure 6C–6J**). Together, these findings demonstrate that sustained G_s_-coupled metabotropic signaling can both prevent and reverse the cellular adaptations that render dSPNs abnormally susceptible to DA-evoked hyperactivation following severe loss of SNc axon innervation.

## Discussion

Although disease severity is an important determinant of the motor effects of DA replacement therapy in PD ^13,24,25,81^, precisely when and how SNc degeneration modifies the responsiveness of striatal circuits to DA remains incompletely understood. Here, we demonstrate that dSPNs transition from physiological to pathological DA responsiveness only following complete local loss of DA axons and identify the prolonged absence of D1 receptor-mediated signaling in dSPNs as a key cell-autonomous mechanism underlying this transition. Our findings establish sustained signaling by G_s_-coupled metabotropic receptors as critical for maintaining dSPNs’ physiological responsiveness to DA and suggest that preserving or restoring this signal in advanced PD may offer a strategy for preventing or reversing maladaptive plasticity contributing to levodopa-evoked motor complications.

Our study builds on established models of DA denervation while incorporating key modifications that address longstanding challenges in isolating the specific contributions of progressive DA axon loss. First, experiments using chemically-induced DA neuron lesions in rodents are typically designed to compare subjects in which DA axons are either intact or severely lesioned, making it difficult to determine precisely when cellular and molecular phenotypes emerge during the progression of PD. Moreover, denervation extent is typically estimated macroscopically across large striatal regions and reported as a singular value (e.g., 90%), which may not accurately reflect the local DA innervation experienced by individual DA-recipient neurons, as DA axon loss is rarely homogeneous throughout the striatum ^3,82,83^. We address these limitations by generating spatial gradients of DA axon denervation within the same subject and by quantifying local DA innervation at cellular resolution. The restricted anatomical extent of lesions our provides a powerful mechanistic approach for dissecting how DA axon loss locally alters striatal microcircuits.

A second challenge is distinguishing plasticity caused by DA denervation from that induced by repeated DA treatment. Although chronic levodopa paradigms appropriately model the cumulative adaptations induced by repeated DA receptor stimulation during PD treatment (commonly referred to as sensitization or priming), they cannot distinguish changes that arise from the persistent loss of DA from those subsequently imposed by repeated stimulation of DA receptors. We therefore used a single, first-ever DA replacement challenge to probe how prior denervation alters the responsiveness of striatal microcircuits to DA, before repeated DA exposure could itself remodel striatal microcircuits. This approach allowed us to isolate functional adaptations that arise at different levels of DA axon loss.

Consistent with previous work ^50,69,70^, we found that dSPNs exhibit the hypoactivity characteristic of parkinsonism in regions with substantial denervation, but respond aberrantly to levodopa in regions devoid of DA axons. Remarkably, this aberrant response did not emerge in partially denervated regions, even when residual DA axon density was exceedingly low. Instead, levodopa restored dSPN activity toward physiological levels, suggesting that the therapeutic response to DA replacement is preserved as long as some DA innervation remains. Interpreting this sharp transition is complicated by the fact that levodopa must first be converted to DA, whose extracellular availability is tightly regulated by DAT-expressing axons. Thus, the spatial difference in dSPN responsiveness could reflect presynaptic differences in DA handling: levodopa-derived DA may be efficiently buffered through DAT-mediated reuptake in regions where residual DA axons remain, but accumulate excessively in regions devoid of DA axons ^32^. To distinguish this potential presynaptic contribution from postsynaptic changes in DA responsiveness, we directly stimulated D1 receptors pharmacologically. This manipulation reproduced the spatial pattern of levodopa-induced c-Fos and Arc expression, indicating that the sharp difference in dSPN responses on either side of the denervation boundary is not primarily attributable to differences in extracellular DA dynamics. Rather, our findings point to a profound postsynaptic change in how dSPNs respond to DA following local DA axon loss.

The marked induction of both c-Fos and Arc in dSPNs following direct D1 receptor stimulation in our model closely recapitulates molecular phenotypes reported in previous DA denervation studies, which attributed their findings to a supersensitivity of dSPNs to DA ^34,84^. While this correspondence provides independent support for the validity of our model, we suggest that the term ‘supersensitivity’ may not fully capture the nature of the adaptation that dSPNs undergo when DA axons are lost, as it implies that D1 receptor signaling is merely amplified. However, our present and previous findings suggest that DA denervation fundamentally alters the manner in which dSPNs respond to D1 receptor signaling. In the DA-intact brain, D1 receptor stimulation does not notably elevate dSPN activity; dSPNs instead follow an inverted-U relationship with DA signaling, approaching maximal activity at physiological DA levels and declining activity when DA signaling is pharmacologically reduced or elevated ^48,49,85^. Following DA depletion, this relationship breaks down, and D1 receptor stimulation instead evokes aberrant dSPN hyperactivation ^48–50^. In line with this, intracellular signaling cascades recruited by D1 receptors change when denervation is severe ^84^. Together, these observations suggest that DA deprivation profoundly transforms the manner in which dSPNs respond to D1 receptor stimulation, rendering them aberrantly susceptible to hyperactivation, which contributes to the production of excessive movements ^47,50^.

Our denervation gradients also allowed us to establish that, although increased dSPN intrinsic excitability is a robust consequence of DA depletion ^50,69,70^, it does not covary with the profound shift in DA responsiveness observed across denervation boundaries and is therefore insufficient to account for it. Instead, the increase in intrinsic excitability may represent a homeostatic response to the hypoactivity that accompanies milder decreases in DA innervation earlier in the progression of PD. The later emergence of a pathological response to DA therefore reflects a distinct form of denervation-induced plasticity, potentially involving changes in synaptic transmission and/or intracellular signaling that remain to be determined ^14^.

What molecular pathways trigger the pathological transformation of dSPNs following DA denervation? The close spatial correspondence between the denervation boundary and the pathological shift in DA responsiveness suggests that plasticity is governed by signals provided by DA axons. Indeed, our genetic and chemogenetic experiments demonstrate that a key trigger for the phenotypic transformation of dSPNs is the prolonged loss of D1 receptor signaling in dSPNs themselves, as (1) deleting D1 receptors sparsely from adult dSPNs reproduced the aberrant response to G_s_-coupled receptor stimulation despite neighboring DA axons and striatal circuitry remaining intact, and (2) sustaining G_s_-coupled metabotropic signaling in dSPNs prevented the emergence of maladaptive plasticity despite complete DA axon loss. Together, these complementary gain-and loss-of-function experiments demonstrate that dSPNs can detect and respond to DA denervation cell-autonomously via D1 receptors and that persistent signaling by metabotropic receptors coupled to the G_s_ family of heterotrimeric G proteins in dSPNs is critical for preventing plastic changes that leave these cells aberrantly susceptible to hyperactivation by DA. Consistent with this, D1 receptors are the predominant G_s_-coupled metabotropic receptors expressed by dSPNs ^86^, making D1 receptor signaling a major source of G_s_ pathway activation in these neurons. These observations therefore suggest that D1 receptors serve dual roles in dSPNs: they acutely regulate dSPN responses to axonal release of DA, while also chronically maintaining the molecular state that constrains those responses.

Our findings have important implications for understanding the consequences of neuromodulatory loss. First, they demonstrate that the loss of a neuromodulatory input can fundamentally transform circuit function, including how neurons respond when that same neuromodulator is reintroduced. This is particularly problematic when interpreting studies that use DA neuron lesions to infer the physiological functions of DA: complete DA denervation does not simply reproduce a state of reduced dopaminergic signaling, it creates a fundamentally altered, pathological circuit whose responses may no longer reflect those of the intact brain.

More broadly, our findings raise the possibility that the denervation-induced dSPN transformation we characterize here determines when DA treatments first transition from being therapeutic to evoking dyskinesia during the progression of PD. Indeed, strong activation of dSPN ensembles is sufficient to evoke dyskinesia^50–55^, yet under normal conditions, D1 receptor stimulation does not produce widespread dSPN hyperactivation (see **Figure 2** at 0–99% denervation levels) ^35,48,49,85^. Our findings and others show that this physiological constraint is lost when dSPNs lose their surrounding DA innervation, leaving them prone to DA-evoked hyperactivity. DA’s ability to produce visible dyskinesia may therefore depend on the extent of striatal regions entirely devoid of DA axons, as pathological hyperactivation of sufficiently large dSPN ensembles is required to produce motor phenotypes ^25,87^ (see Limitations below). This framework raises the possibility that the pathological denervation-related transition of dSPN responsiveness to DA contributes to the relationship between the severity of PD (or time since PD diagnosis) and the emergence of dyskinesia, including the observation that limbs most affected by the disease are often the first to develop motor complications ^4,15–17^. In this view, the initial emergence of dyskinesia may mark a natural stage in the progression of PD, arising when DA denervation has progressed sufficiently for large ensembles of dSPNs to respond aberrantly to DA replacement. Once dSPNs have entered this pathological state, additional factors – including treatment dose, extracellular DA dynamics, and plastic changes in dSPNs and other striatal populations following repeated DA exposure – may shape how dyskinetic behavior is ultimately expressed, including its intensity, duration and temporal pattern.

Lastly, the ability of sustained G_s_-coupled metabotropic signaling to both prevent and reverse the denervation-induced pathological transformation of dSPNs suggests that this state is not irreversible and, perhaps most encouragingly, that it may be leveraged therapeutically to either reduce the severity of dyskinetic episodes or prevent their emergence outright. In patients with PD, this might be approached by minimizing prolonged periods of DA withdrawal. Recent clinical data indicate that continuous delivery of levodopa effectively reduces troublesome dyskinesia in patients with advanced PD ^88–90^. Whereas the efficacy of continuous treatments is attributed to their ability to avoid excessive DA receptor stimulation, our results suggest that sustaining a minimal level of dopaminergic signaling in dSPNs may be equally important to prevent dSPNs from becoming abnormally vulnerable to DA-mediated hyperactivation. Longer-acting or continuous dopaminergic therapies may therefore reduce the severity of dyskinesia by blunting levodopa peaks as well as preventing or reversing the maladaptive plasticity that dSPNs undergo when degeneration becomes complete. Previous work in a developmental model of DA deficiency lends support to this view ^91^. Future experiments will test this hypothesis in established models of dyskinesia.

Together, our findings suggest that an important form of striatal maladaptive plasticity implicated in DA replacement-induced dyskinesia emerges when local DA axon loss becomes complete, late in the progression of PD. Elucidating the molecular and cellular mechanisms governing this transition during disease progression may reveal new strategies for preventing or reversing these maladaptive changes while preserving the therapeutic benefits of DA replacement.

## Limitations

Several limitations of our study warrant consideration. First, although our sparse cellular manipulations enabled us to investigate denervation-induced plasticity while minimizing potential secondary effects of dyskinetic movements, they did not target a sufficiently large ensemble of dSPNs to assess whether preventing or reversing this plasticity alters levodopa-induced dyskinesia ^25^. Thus, whether the cellular adaptations identified here are sufficient to drive motor complications, and whether their prevention or reversal can mitigate dyskinesia remain open questions. Future studies targeting larger dSPN ensembles will be needed to establish the behavioral consequences of these manipulations. Second, our assessment of the pathological response to DA relied primarily on the activity-associated immediate early gene products c-Fos and Arc. Although their complementary expression patterns provide convergent evidence for denervation-dependent changes in dSPN activation, neither directly measures neuronal activity. Functional measurements, such as *in vivo* electrophysiological recordings or calcium imaging will be important to determine how gain-and loss-of-function manipulations alter dSPN activity dynamics and to further establish the relationship between the molecular adaptations identified here and the pathological neuronal responses to DA observed *in vivo*. Finally, our approach modeled different degrees of DA axon denervation across space rather than gradually over time, as in PD. Whether dSPNs undergo similar changes in DA responsiveness as denervation gradually worsens remains to be determined. This question could be addressed in models of progressive nigrostriatal degeneration, such as MitoPark mice ^24^, by relating local DA axon density to dSPN responses to a first DA challenge at different stages of the disease.

## Acknowledgements

We thank members of the Tritsch laboratory for comments on the manuscript, and Tim Cheung and Yunmin Ding in the Kang laboratory for helpful advice on the experiments. We acknowledge the Neuroscience Institute’s imaging facilities for microscope availability and the Douglas’ Molecular and Cellular Microscopy Platform. This work was supported by NIH 5TL1TR001447 and 5T32NS086750 (JRM); McGill Conrad F. Harrington Fellowship and CIHR CPRA (SP); NIH 5R35NS116858 (AEC and JSD), R01MH130658 (TS) and R21NS133640 (UJK and NXT); the Parkinson’s Foundation (NXT); the Canada First Research Excellence Fund, awarded through the Healthy Brains, Healthy Lives initiative at McGill (NXT); CIHR Project Grants 205940 and 213008 (NXT); and FRQ Chercheur Boursier octroi 365987 (NXT). The funders had no role in study design, data collection and analysis, decision to publish or preparation of the manuscript.

## Author Contributions Statement

JRM, MM and NXT conceived of the project. JRM, UJK and NXT designed experiments, interpreted data, and wrote the manuscript with inputs from all authors. JRM performed and analyzed histological experiments, with initial help from MM. SP, MM and RM performed and analyzed electrophysiology experiments. AEC helped with histology, imaging and quantification. JSD, TS, UJK and NXT provided supervision and funding.

## Methods

### Animals

All procedures were performed in accordance with protocols approved by the NYU Langone Health Institutional Animal Care and Use Committee (protocol 170123) and the Douglas Research Centre Animal Use and Care Committee (protocol 10058). Mice were housed in a reverse 12-hr light-dark cycle with *ad libitum* access to food and water. The following strains available from The Jackson Laboratory were used for experiments and maintained on a C57Bl6/J (Jackson Laboratory; 000664) background: heterozygous *Drd1-tdTomato* transgenic mice (Jackson Laboratory; 016204), homozygous *Drd1*^flox/flox^ mice (Jackson Laboratory; 025700) and heterozygous *Drd1-Cre* transgenic mice (EY217 line; Gensat; #030778-UCD). Adult mice (8+ weeks) of both sexes were used for experiments. We did not observe differences in any of the reported phenotypes between male and female mice; results from both sexes were therefore pooled in figures and for analyses.

### Adeno-associated viruses (AAVs)

To permit the identification of rM3Ds-expressing dSPNs, we generated a Cre-dependent bicistronic AAV under control of the EF1α promoter that encodes the G_s_-coupled Designer Receptors Exclusively Activated by Designer Drugs (DREADD) rM3Ds ^79^ tagged on the 3’ end with three FLAG sequences along with the nuclear marker histone 2B (H2B) fused to three HA tags, both separated a by P2A sequence (AAV-EF1α-DIO-rM3Ds-3xFLAG-P2A-H2B-3xHA; referred to as AAV-DIO-rM3Ds-P2A-H2B in figures and text). The expression plasmid (to be deposited at Addgene) was custom synthetized (Genscript), maxi-prepped and packaged into AAV5 and AAV8 (WZ Biosciences/AAVnerGene Inc.; titer: 1.2 × 10^13^ gc/mL). AAV5 provided the best expression in dSPNs. All other AAVs were obtained from Addgene and used undiluted: AAVrg-pgk-Cre (#24593; 1.4 × 10^12^ gc/mL) and AAV1.CMV.PI.Cre.rBG (#105537; 1.2 × 10^13^ gc/mL).

### Stereotaxic surgery

Intracranial delivery of AAVs and 6-hydroxydopamine (6OHDA) were performed using standard techniques, as before ^48,92^. Briefly, mice were anesthetized with isoflurane, placed in a stereotaxic apparatus (KOPF Model 1900) on a heating pad (Harvard Apparatus) and administered ketoprofen (10 mg/kg in 0.9% saline) and 0.9% saline (up to 1 mL) subcutaneously. Mice were also given Desipramine (25 mg/kg, i.p.) 30 minutes prior to intracranial 6OHDA infusions. After exposing and cleaning the skull under aseptic conditions, craniotomies were made with a drill fitted with a ball-shaped (0.4 mm diameter) carbide bur (Foredom). Intracranial injections were performed using a Nanoject III (Drummond; rate: 50-120 nl/min) fitted with a pulled micropipette (Drummond; Wiretrol II Calibrated Micropipets) back-filled with mineral oil and front-filled with AAV, 6OHDA or vehicle. Gradients of DA axons were created by infusing 1 µl 6OHDA (5 µg/µl in 0.9% saline and 0.2% ascorbic acid) in dorsolateral striatum (DLS; 0.2 AP, 2.4 ML,-2.7 DV; all coordinates in mm, from bregma and pial surface). Sham lesions were performed at the same coordinates on the contralateral side using vehicle (1 µl 0.9% saline and 0.2% ascorbic acid). In Figure S2, gradients centered on the dorsomedial striatum used coordinates 0.7 AP, 1.3 ML and-2.1 DV, and medial forebrain bundle lesion used-1.3 AP, 1.2 ML and-5.4 DV. To retrogradely transduce dSPNs, 200 nl AAVrg-pgk-Cre was injected in substantia nigra pars reticulata (SNr;-3.3 AP, 1.5 ML,-4.5 DV) of *Drd1*^flox/flox^ or *Drd1*^wt/wt^ littermates. To express rM3Ds in dSPNs, 400-600 nl of AAV-DIO-rM3Ds-P2A-H2B (serotype 5 or 8) was infused in DLS (AP: 0.6; ML: 2.3; DV:-2.5). Injection needles were left in place for 5-10 min prior to slowly exiting the brain. Mice receiving 6OHDA injections were given 1.0 mL 5% glucose in 0.9% saline after surgery. Mice received three days of post-op care consisting of Ketoprofen (10 mg/kg, s.c.) or Ketoprofen and 5% glucose in 0.9% saline if surgery included 6-OHDA lesions. Animals receiving DCZ water prior to 6OHDA lesioning (see **Figure 5**) instead received Ketoprofen and a combination of 5% glucose and 10 mg/L DCZ in 0.9% saline (1 ml, s.c.) on the day of the surgery, and for three days after for post-op care.

### Drugs

The following drugs were administered intraperitoneally such that each mouse received 0.1 mL solution per 10 g body weight: levodopa methyl ester hydrochloride (Tocris; Cat# 3788; 5 mg/kg) co-administered with benserazide hydrochloride (Millipore; #B7283; 12 mg/kg) in sterile 0.9% saline; SKF81297 hydrobromide (Tocris; #1447; 3 mg/kg) in sterile 0.9% saline; and deschloroclozapine dihydrochloride (DCZ; Hello Bio; #HB9126; 200 μg/kg) in pico-pure water. Animals receiving DCZ in the drinking water (10 mg/L) via regular cage bottle had *ad-lib* access, with a fresh batch made every 4 days.

### Immunohistochemistry

Unless noted otherwise, mice were perfused 4 weeks after 6OHDA lesions and 90 min after acute challenge with levodopa, SKF81297, DCZ or saline to probe the responsiveness of dSPNs to G_s_-coupled metabotropic receptor signaling. Animals were deeply anesthetized with isoflurane and transcardially perfused with phosphate buffered solution (PBS) followed by 4% paraformaldehyde (PFA) in PBS. Brains were post-fixed in 4% PFA overnight at 4°C, washed twice with PBS, and transferred to PBS for storage at 4°C until sliced and immunostained. Free-floating 50 μm coronal sections were obtained using a vibratome (Leica; VT1000S or VT1200S) and permeabilized and blocked with 5% normal goat serum (NGS; Abcam; ab7481) in PBS supplemented with 0.1% Triton-X-100 (PBS-T; Sigma; T9284) for 1 hour at room temperature. Sections were then stained with the following primary antibodies in 1% NGS in PBS-T overnight at room temperature: NeuN (Millipore; MAB377; 1:500), c-Fos (Synaptic Systems; 226 003; 1:1000), dopamine transporter (DAT; Millipore; MAB369; 1:1000), HA (Abcam; ab18181 at 1:500 or ab9110 at 1:1000, or ab9111 at 1:500), Arc (Synaptic Systems; 156 005; 1:500), Cre (Synaptic Systems; 257 005; 1:500), and/or D1 dopamine receptor (Sigma-Aldrich; D2944; 1:500). The following secondary antibodies were applied for 1–2 h at room temperature in 1% NGS in PBS-T (all at 1:500 dilution unless otherwise noted): donkey anti-chicken IgY/IgG Cy3 (Jackson ImmunoResearch; 703-165-155, at 1:1000), goat anti-mouse IgG Alexa Fluor Plus 405 (Thermo Fisher Scientific; A48255), goat anti-rabbit IgG Alexa Fluor 488 (Thermo Fisher Scientific; A-11034), goat anti-mouse IgG Alexa Fluor 568 (Thermo Fisher Scientific; A-11031), goat anti-rat IgG Alexa Fluor 647 (Thermo Fisher Scientific; A-21247), and/or goat anti-guinea pig IgG Alexa Fluor 647 (Thermo Fisher Scientific; A-21450). Brain sections were mounted on Superfrost plus slides (Fisher; 12-550-15) and coversliped (Fisher; 12-541-035) with Fluoromount-G mounting medium without DAPI (SouthernBiotech; 0100-01) or DAPI Fluoromount-G (SouthernBiotech; 0100-20). Immunofluorescence images were obtained first on an Olympus VS120 slide-scanning microscope using a 10X objective before acquiring higher resolution confocal images with a 40X objective on either a Zeiss LSM700 or LSM800, or Olympus FV1200 confocal microscopes. For all experiments, the right and left hemispheres were both acquired at the same time using identical parameters. In figures, fluorescence signal in individual channels is shown separately as grayscale (with color-coded headings), and in color in merged image.

### Slice Electrophysiology

Acute brain slices were obtained from adult mice (N = 20; median age: 89 days old) as before ^73^. Mice were anaesthetized by isoflurane inhalation and perfused intracardially with oxygenated ice-cold choline-based cutting solution containing (in mM) 110 choline chloride, 25 NaHCO3, 2.5 KCl, 7 MgCl2, 0.5 CaCl2, 1.25 NaH2PO4, 25 glucose, 11.6 ascorbic acid, and 3.1 pyruvic acid. The brain was removed, blocked, and transferred to a slicing chamber, all while submerged in oxygenated ice-cold cutting solution. Coronal slices of striatum were cut (thickness: 270 μm) using a Leica VT1000s vibratome, transferred for 10-20 min to a holding chamber submerged in artificial cerebrospinal fluid (ACSF) containing (in mM) 125 NaCl, 2.5 KCl, 25 NaHCO3, 2 CaCl2, 1 MgCl2, 1.25 NaH2PO4, and 11 D-glucose, at 34°C and subsequently maintained at room temperature (21-22°C) until use. All recordings were obtained within 4 hours of slicing. Both cutting solution and ACSF were constantly oxygenated with 95% O_2_/ 5% CO_2_.

Individual slices were transferred to a recording chamber mounted on an upright microscope (Slicescope; Scientifica) and continuously superfused (2–3 mL per minute) with ACSF warmed to 32-34°C by an in-line heater (SH-27B; Warner Instruments). Cells were visualized through a 40X water-immersion objective with infrared differential interference contrast optics. dSPNS were identified based on the presence of tdTomato under epifluorescence illumination as well as their firing properties. Patch pipettes (2–4 MΩ) pulled from borosilicated glass (G150F-4, Warner Instruments) and were filled with internal solution containing (in mM) 135 KMeSO_3_, 3 KCl, 10 HEPES, 1 EGTA, 0.1 CaCl_2_, 4 Mg-ATP, 0.3 Na-GTP, 8 Na_2_-Phosphocreatine (pH 7.3 adjusted with KOH; 295 mOsm·kg^−1^). To assess intrinsic excitability, depolarizing current steps (in 50 pA increments) were applied at regular intervals and the resulting rate of action potentials fired was quantified as spikes per second. dSPNS expressing tdTomato were identified using epifluorescence.

### Quantification and Statistical Analysis

Immunofluorescence images were analyzed using FIJI (ImageJ2 version 2.16.0/1.54p) and custom code to be deposited to GitHub. Data are reported in text and figures as mean ± s.e.m. Statistical tests were run in Prism 11 (Graphpad) using non-parametric Mann-Whitney U test for independent groups of mice, Wilcoxon signed rank test for paired data, nested t test for randomly subsampled cells and Kruskal-Wallis one-way ANOVA (for balanced data) or mixed-effects analysis (for unbalanced data) with *post-hoc* Dunnett multiple comparison tests for comparisons between multiple groups, as indicated in figure legends. *N* denotes the number of mice, and *n* dSPNs, unless stated otherwise. *p* values smaller than 0.05 were considered statistically significant and assigned the following nomenclature in figures: \**p* < 0.05, \*\**p* < 0.01, \*\*\**p* < 0.005 and \*\*\*\**p* < 0.001.

To determine the local density of DA axons as shown in Figure 1, we took confocal images of DAT staining throughout the entire dorsal striatum in the right and left hemisphere and randomly seeded 2500 circular regions of interest (ROIs; 50 µm diameter each) throughout each. The center of each ROI (*x,y* coordinates) was used to assign their position along the medio-lateral and dorso-ventral axes of the striatum. For each ROI, DAT immunofluorescence was binarized to highlight DA axon processes, and the fraction of neuropil occupied by DA axons was calculated to obtain a measure of axonal density that is not directly related to DAT fluorescence intensity. Thresholds used for binarization of DAT signals were determined empirically for each mouse from the 6OHDA-lesioned hemisphere and set to capture continuous, fiber-like DA axon processes while avoiding isolated pixels within the neuropil, which are unlikely to reflect DA axons. The same threshold was then applied to the sham-lesioned hemisphere. In Figure 1D, ROIs were grouped in 100-µm bins along the medio-lateral axis and, for each bin, DA axon density measures were averaged and normalized to the mean DA axon density of the 5 medial-most bins (i.e. dorsomedial striatum). For each lesioned striatum, we manually outlined a DA axon denervation boundary from raw confocal images under high contrast to identify the furthest extent that fiber-like DAT^+^ axonal processes were observed along the medio-lateral axis before entering zones devoid of axons in lateral-most striatum. The same denervation boundary was then applied to the contralateral (sham-lesioned) striatum for quantification purposes. The percent DA axon denervation in Figure 1F was obtained by finding the smallest distance separating each ROI to the DA denervation boundary in sham-and 6OHDA-lesioned striatum, by averaging DA axon density across ROIs (in 100-µm bins relative to denervation boundary) and by applying the following calculation for each bin: 100% – [(density_sham_ – density_6OHDA_) * 100 / density_sham_]. To estimate the density of DA axons that individual dSPNs are locally exposed to (Figure 2), we extracted DA axon degeneration values from spatial maps using the (*x,y*) coordinates of each imaged dSPNs.

To quantify the expression of molecular markers in dSPNs, we first obtained the outline (i.e. mask) of dSPNs by applying the Cellpose segmentation algorithm to either the endogenous somatic fluorescence signal in *Drd1-tdTomato* mice (Figure 2) or to nuclear H2B immunofluorescence (revealed with HA tag) in Cre^+^ dSPNs (Figures 4, 5 and 6). Masks were checked for accuracy and, when necessary, redrawn manually. For each dSPN mask, the immunofluorescence intensity of c-Fos and Arc was measured and normalized to a mean background fluorescence value obtained from 5 neuropil measurements within the same hemisphere (for c-Fos) or contralaterally (for Arc). dSPNs were deemed positive for c-Fos if background-normalized fluorescence exceeded a threshold of 3. dSPNs were deemed positive for Arc if background-normalized fluorescence exceeded a threshold of 2, except for Figure 4, where a more stringent threshold of 4 was used. The intensity of c-Fos and Arc signal in positive cells was only reported if at least 5 cells were deemed positive. Across experiments, we report mean values obtained for all imaged dSPNs per experimental subject (*N =* 3-8 subjects per experimental condition). In some figures, we also plot values obtained in individual neurons (*n*) across subjects; to avoid biases stemming from uneven sampling, the same number of cells was drawn randomly from each experimental subject. In Figures 5 and 6, only rMD3s^+^ dSPNs within DA-depleted zones (confirmed to be at minimum 100 µm from DA axon boundary) were quantified.

The functional imaging dataset from Maltese *et al* ^48^ was re-analyzed in Figure S2 by distinguishing mice in which 6OHDA-mediated lesioning of SNc axons led to partial or complete loss of striatal DA axons using epifluorescence images of fixed tissue immunostained for DAT. We previously reported that 6OHDA decreased DAT immunolabeling in dorsal striatum by 91 ± 3% (*N =* 12 mice; range: 70–100 %) compared to sham controls using pixel intensity measures obtained macroscopically across the entire dorsal striatum. In light of our c-Fos and Arc findings, we re-examined these sections at high magnification, distinguishing mice in which some DA axons remained visible in regions imaged functionally (i.e., 70-99% denervation; *N* = 7 mice) vs. not (i.e., 100% denervation; *N* = 5 mice). For each group, we then re-plotted the size of locomotion-activated dSPN ensembles imaged longitudinally across the same fields of view (1-2 per mouse) using the Ca^2+^ indicator GCaMP6f at 3 experimental time points: before lesioning nigrostriatal DA neurons with 6OHDA, 30 days after the lesion and following acute challenge with levodopa (10 mg/kg along with 12 mg/kg benserazide hydrochloride, i.p.). Activity measures per field of view were normalized to pre-lesion values and averaged per mouse.

---

**Figure S1.**
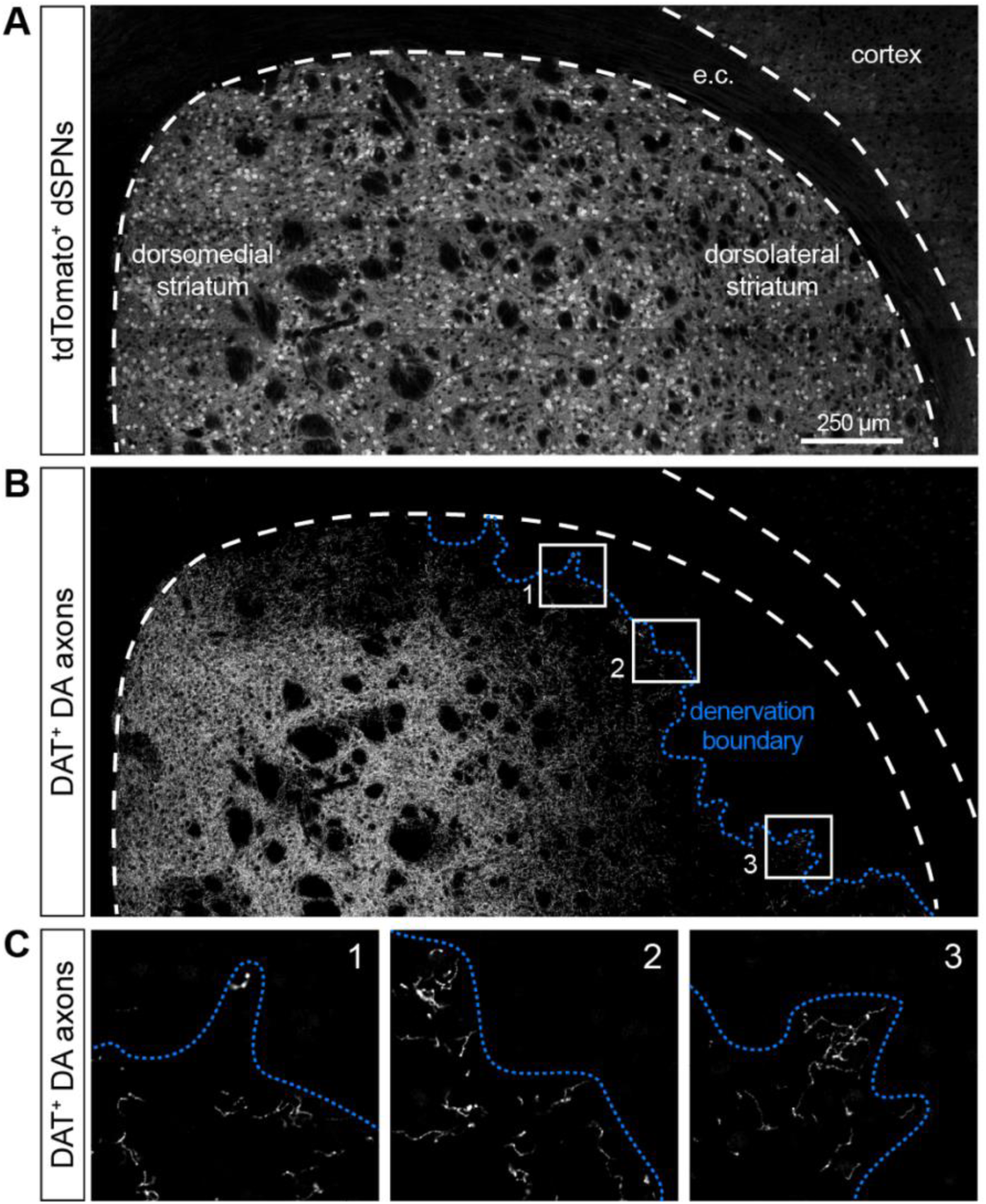
(related to Figure 1): Example denervation boundary along 6OHDA gradient. Grayscale confocal image of dorsal striatum showing tdTomato^+^ dSPNs (A) and DA axons immunostained for DAT (B). Dotted blue line in (B) depicts the DA axon denervation boundary, which we define as the furthest extent that DAT^+^ axonal processes are observed as one moves down the DA denervation gradient (i.e., toward the center of the 6OHDA infusion) before entering zones devoid of axons in dorsolateral striatum. White dashes outline striatum and cortex, separated by the external capsule (e.c.). Labeled white boxes in (B) are shown at higher magnification in (C) to further illustrate how the denervation boundaries are drawn relative to DA axons.

**Figure S2.**
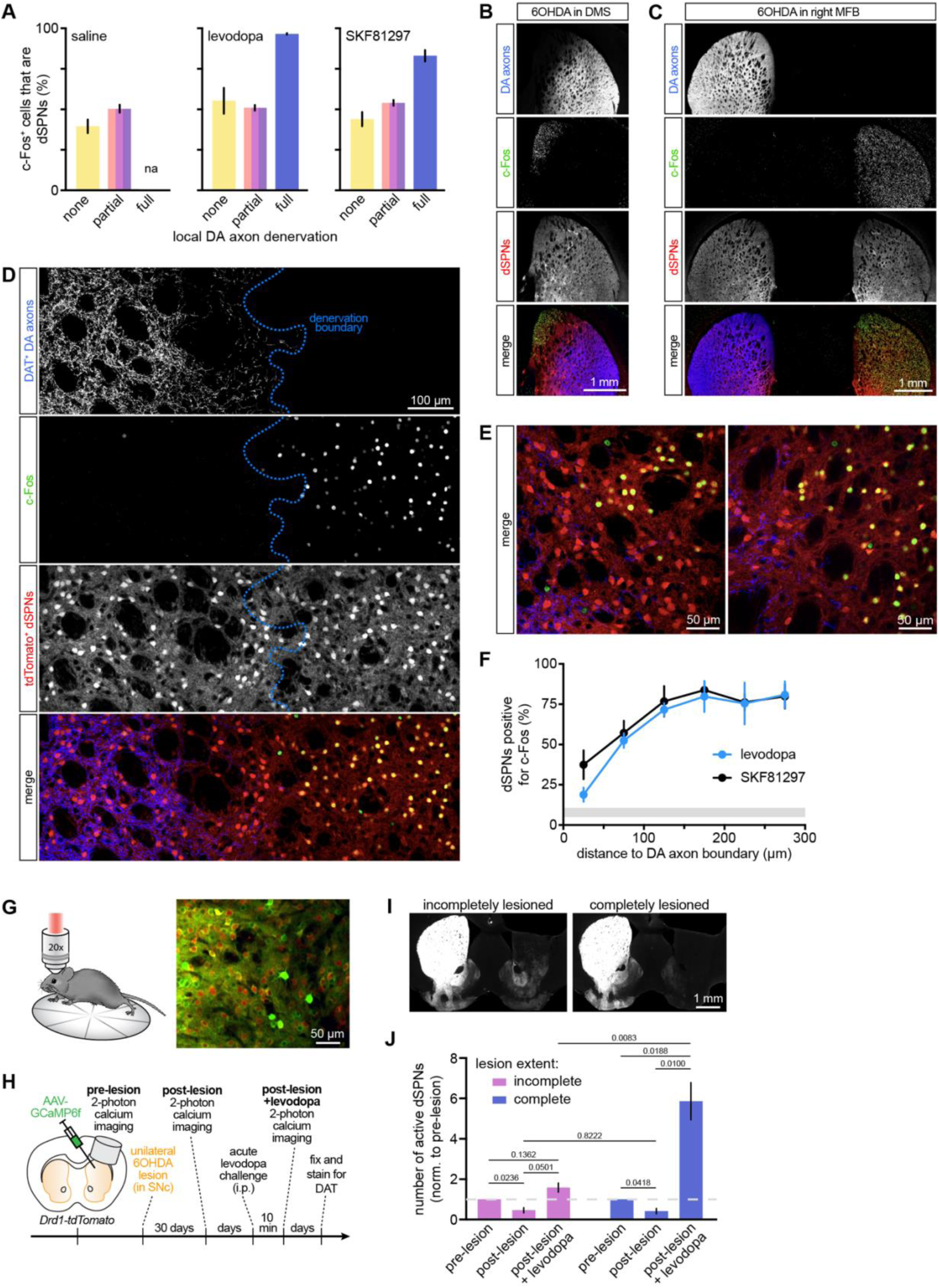
(related to Figure 2): dSPNs become aberrantly prone to DA-mediated hyperactivation when they lose local DA innervation. A. Mean (± s.e.m.) percentage of striatal c-Fos^+^ nuclei that are tdTomato^+^ dSPNs in mice treated with saline (*N =* 7), levodopa (*N =* 5) or SKF81297 (*N =* 7), binned according to local DA axon denervation extent. B. Coronal image of dorsal striatum in a *Drd1-tdTomato* mouse with 6OHDA gradient centered on dorsomedial striatum (DMS) following first-ever challenge with levodopa. Note the selective induction of c-Fos in dorsomedial-most striatal regions devoid of DA axons immunolabeled for DAT. C. Same as (B) for a *Drd1-tdTomato* mouse with 6OHDA infused into the right medial forebrain bundle (MFB), producing complete loss of DA axons throughout the dorsal striatum and corresponding striatum-wide induction of c-Fos following levodopa challenge. D. Confocal image of DAT, c-Fos and tdTomato fluorescence (individual channels shown as gray scale, merged in color) in striatal region straddling the DA axon denervation boundary (*dotted blue line*) following first-ever challenge with levodopa. Note the strong induction of c-Fos (the vast majority of which are dSPNs) on the right side of the denervation boundary, where DA axon loss is complete. E. Same as (D) for two other tdTomato mice challenged with levodopa. The DA denervation boundary is not drawn for clarity; surviving DA axons (*blue*) occupy the left half of each image. dSPNs respond aberrantly to levodopa in regions devoid of DA axons, as evidenced by strong induction of c-Fos (*green*). F. Mean (± s.e.m.) percentage of dSPNs that are c-Fos^+^ in mice challenged with levodopa (*blue*; *N =* 5) or SKF81297 (black; *N =* 7), binned according to the shortest distance of individual dSPNs to the DA axon denervation boundary. Gray zone depicts the percentage of dSPNs that are normally c-Fos^+^ in striatal regions where DA axons are intact. Note that dSPNs respond similarly to levodopa or direct stimulation of D1 receptors, pointing to a ‘post-synaptic’ locus of maladaptive plasticity that is strongly dependent on physical distance to the nearest surviving DA axon. G–J. New analysis of functional Ca^2+^ imaging dataset from Maltese et al. 2021 ^48^. G. Schematic of experimental setup (*left*), in which the dorsal striatum of *Drd1-tdTomato* mice expressing the Ca^2+^ indicator GCaMP6f in all striatal neurons was imaged by 2-photon microscopy through an implanted imaging window while mice locomote on a circular motorized treadmill imposing locomotor bouts of constant speed and duration. *Right*, maximum projection image showing dSPNs (*red*) and GCaMP6f-expressing striatal neurons (*green*). H. Experimental timeline. GCaMP6f was expressed virally in all striatal neurons using a synapsin promoter. The activity of tdTomato^+^ dSPNs was monitored during locomotor bouts before and after lesioning midbrain DA neurons with 6OHDA, and shortly after levodopa challenge. I. Coronal forebrain sections immunostained for DAT were re-examined to determine the severity of DA axon denervation in imaged dorsal striatal regions. Although all lesioned hemispheres experienced significant DA axon denervation, some were incomplete (*left*), with some surviving axons visible in dorsal striatum, while others were complete (*right*). To determine how levodopa-evoked dSPN activity varies with DA axon denervation severity, we split our imaged cohort (*N =* 12 mice) into mice where *post-hoc* analyses detected residual DA axons in dorsal striatum (*N =* 7) vs. those where lesions were complete (*N =* 5). J. Mean (± s.e.m.) number of locomotion-related dSPNs activated in dorsal striatum normalized to pre-lesion values in mice with incomplete (*N =* 7) vs. complete (*N =* 5) DA axon denervation. *P* values are indicated in figure (two-way ANOVA; main effects: denervation severity: *p* = 0.0006; imaging time point: *p* < 0.0001; interaction: *p* = 0.0003; *post-hoc* Šídák’s multiple comparisons tests). Note how both groups showed similar dSPN hypoactivation following 6OHDA lesioning of SNc, but differed markedly in how dSPNs respond to levodopa, with dSPNs in incompletely-lesioned dorsal striatum returning to pre-lesion activity levels and dSPNs in fully-lesioned mice reaching activity levels on average 5.9-fold higher (range: 3.9–9.0) than normally observed in DA-intact dorsal striatum.

**Figure S3.**
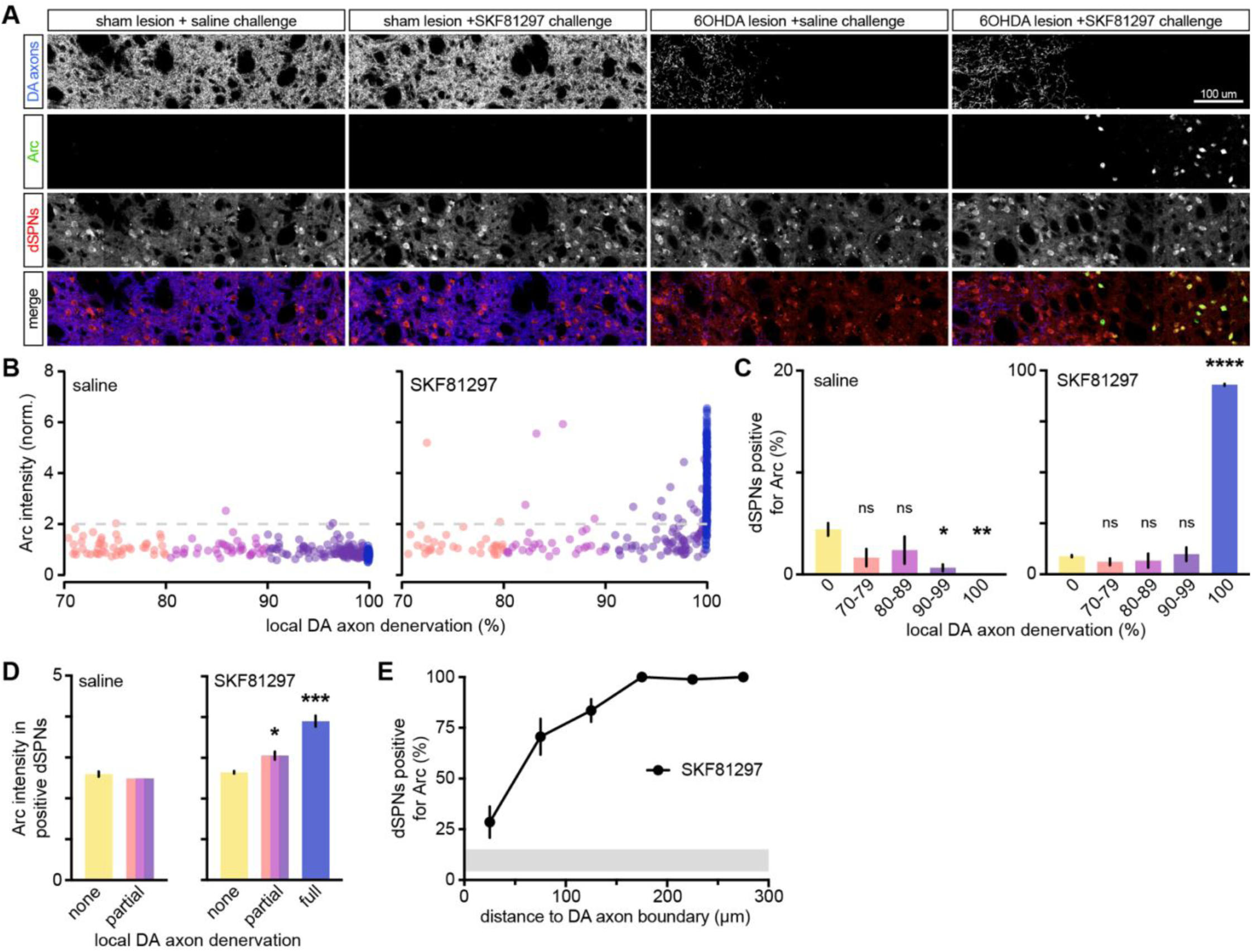
(related to Figure 2): Pathological response of dSPNs to D1 receptor stimulation revealed using Arc immunofluorescence. A. High-magnification confocal images of DAT, Arc and tdTomato fluorescence (individual channels shown as gray scale, merged in color) in *Drd1-tdTomato* mice sham-or 6OHDA-lesioned in dorsolateral striatum shortly after first-ever challenge with saline or SKF81297 (3 mg/kg). See Figure 2A for experimental timeline. SKF81297 only induces Arc expression in dSPNs in regions devoid of DA axons. B. Example intensity of Arc fluorescence (normalized to neuropil) measured in dSPNs exposed to different levels of local DA axon denervation after saline (*left*) or SKF81297 (*right*) challenge. Colors denote DA axon denervation bin (70-79%, 80-89%, 90-99% and 100%). C. Mean (± s.e.m.) percentage of all dSPNs considered positive for Arc at different local DA axon denervation levels following challenge with saline (*N =* 3; one-way ANOVA, *p* = 0.0187; Dunnett’s *post-hoc* tests: 70-79%, *p* = 0.0901; 80-89%, *p* = 0.2533; 90-99%, *p* = 0.0205; 100%, *p* = 0.0078; all vs. 0% denervation), or SKF81297 (*N =* 3; one-way ANOVA, *p* < 0.0001; Dunnett’s *post-hoc* tests: 70-79%, *p* = 0.8102; 80-89%, *p* = 0.8904; 90-99%, *p* = 0.9929; 100%, *p* < 0.0001; all vs. 0% denervation). D. Same as (C) for mean intensity of Arc fluorescence in Arc^+^ dSPNs after challenge with saline or SKF81297 (one-way ANOVA, *p* = 0.0003; Dunnett’s *post-hoc* tests: none vs. partial, *p* = 0.0251; none vs. full, *p* = 0.0002). Too few dSPNs in saline-treated partially-and fully-lesioned striatum were Arc^+^ for statistical analyses. E. Mean (± s.e.m.) percentage of dSPNs that are Arc^+^ in the denervated region of mice challenged with SKF81297, binned according to the shortest distance of individual dSPNs to the DA axon denervation boundary. Gray zone depicts the percentage of dSPNs that are normally Arc^+^ in striatal regions where DA axons are intact.

**Figure S4.**
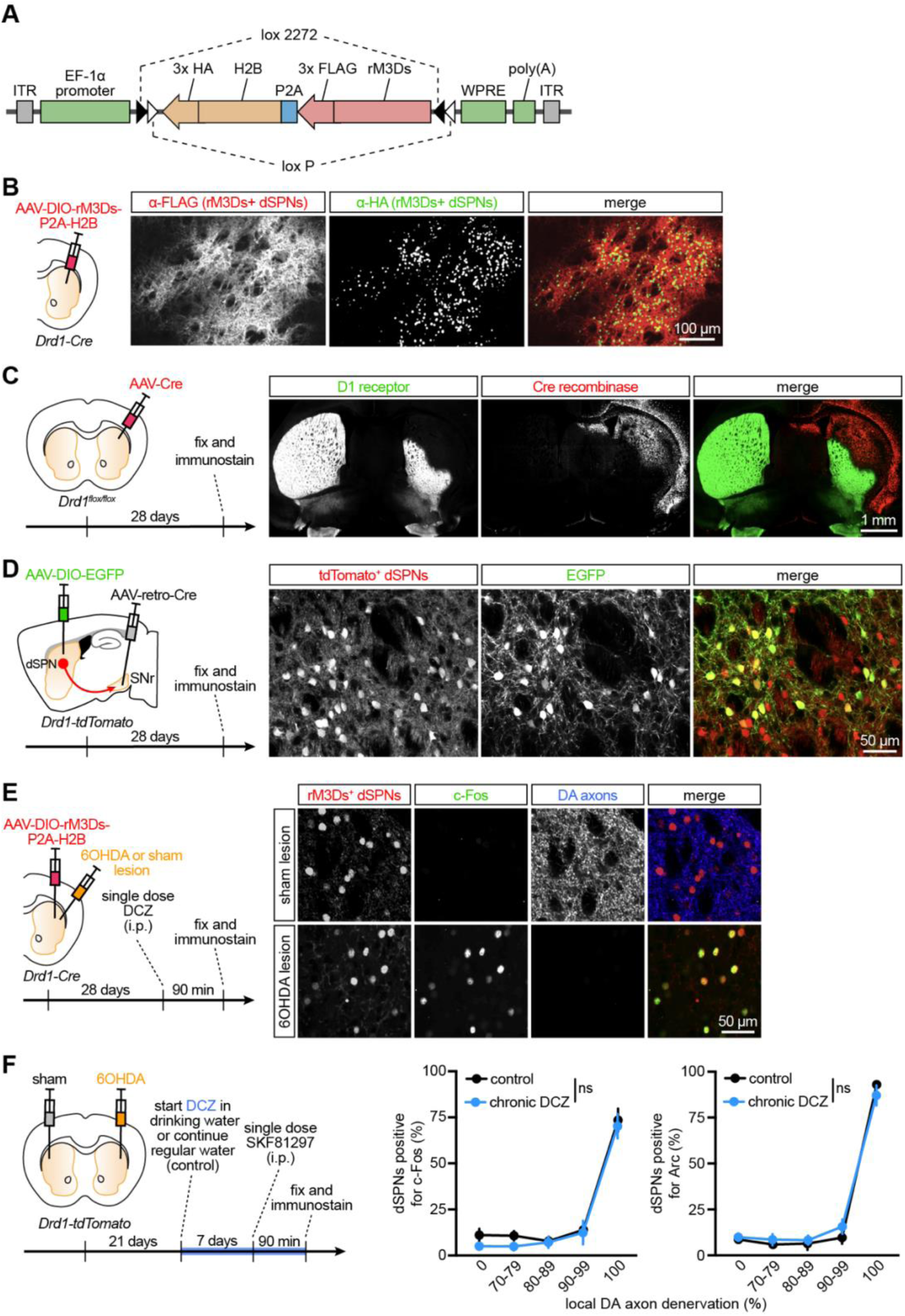
(related to Figure 4): Technical controls for assessing how dSPNs respond to the stimulation of G_s_-coupled metabotropic receptors. A. Schematic of AAV-DIO-rM3Ds-P2A-H2B construct designed to enable bi-cistronic co-expression of the chemogenetic actuator rM3Ds and the nuclear marker H2B in a Cre-dependent fashion. rM3Ds and H2B are respectively tagged with 3 FLAG and HA sequences each to facilitate immunolabeling. DIO: double-floxed open reading frame; the loxP and lox2272 sequences permit Cre-mediated recombination and inversion of rM3Ds/H2B’s open reading frames with respect to the EF-1α promoter. P2A: porcine teschovirus 2A sequence allowing for two separate protein products to be produced from a single mRNA transcript. ITR: inverted terminal repeats. WPRE: woodchuck polyresponse element. Poly(A): human growth hormone polyadenylation. B. Construct validation. *Left*, striatal dSPNs were transduced with AAV-DIO-rM3Ds-P2A-H2B-HA in Drd1-Cre mice. *Right*, epifluorescence image of dorsal striatum region transduced with AAV-DIO-rM3Ds-P2A-H2B, fixed, sectioned in the coronal plane and immunostained using anti-FLAG (*red*) and anti-HA (*green*) antibodies, confirming distribution of rM3Ds in neuropil and H2B in nucleus. C. Validation of AAV-mediated ablation of D1 receptors from striatal neurons. *Left*, AAV-Cre was injected in the dorsolateral striatum of *Drd1*^flox/flox^ mice. *Right*, four weeks after viral transduction, coronal forebrain sections were immunostained using antibodies raised against D1 receptor (green) and Cre recombinase (*red*). Note the absence of D1 receptor protein in striatal regions expressing Cre. D. Validation of retrograde viral approach to selectively target dSPNs from their axon terminals in SNr. *Left*, sagittal diagram of *Drd1-tdTomato* mouse brain showing dSPN cell body (red circle) in striatum and axon (red arrow) projecting to SNr. dSPNs are the only striatal neurons projecting directly to SNr; they can therefore be selectively targeted by injecting into SNr AAVs designed to travel retrogradely from axon terminals to the cell body (AAV-retro-Cre). Retrogradely-targeted Cre^+^ neurons are revealed by injecting another AAV expressing EGFP in a Cre-dependent fashion (AAV-DIO-EGFP). *Right*, confocal image of striatum showing that EGFP is selectively expressed in tdTomato^+^ dSPNs. Note that this retrograde viral approach only labels a sparse subset of dSPNs. E. Confirmation that stimulation of rM3Ds can substitute for D1 receptors to reveal dSPN maladaptive plasticity in DA-depleted mice. *Left*, rM3Ds was virally expressed in dSPNs of *Drd1-Cre mice*. Striatal DA axons were locally lesioned with 6OHDA during the same surgery. Four weeks later, mice were administered a single, first-ever dose of the selective rM3Ds agonist DCZ before being fixed and immunostained for HA (to reveal the nucleus of rM3Ds^+^ dSPNs), c-Fos and DAT. *Right*, example confocal images of rM3Ds^+^ dSPNs in DA-intact (*top*; sham-lesioned) and DA-depleted (*bottom*; 6OHDA-lesioned) immunostained for c-Fos (green) and DAT (blue). Note strong DCZ-mediated induction of c-Fos in rM3Ds^+^ dSPNs in striatal regions devoid of DA axons (i.e., where dSPNs underwent maladaptive plasticity), consistent with previous work ^52^. F. Experimental timeline (*left*) of control experiment showing that chronic administration of DCZ in drinking water does not, on its own, modify denervation-evoked maladaptive plasticity in dSPNs or SKF81297-mediated hyperactivation of locally-denervated dSPNs, as measured with c-Fos (*middle*) or Arc (*right*) immunofluorescence (control cohort on regular water: *N* = 7 for c-Fos, replotted from Figures 2E and *N* = 3 for Arc, replotted from Figure S3C; Chronic DCZ cohort: *N* = 5 mice for both c-Fos and Arc; two-way ANOVA for treatment: *p* = 0.2907 for c-Fos, *p* = 0.8242 for Arc).

## References

1. Kalia LV, and Lang AE (2015) Parkinson’s disease. Lancet 386, 896–912.

2. Burke RE, and O’Malley K (2013) Axon degeneration in Parkinson’s disease. Exp Neurol 246, 72–83.

3. Kordower JH, Olanow CW, Dodiya HB, Chu Y, Beach TG, Adler CH, Halliday GM, and Bartus RT (2013) Disease duration and the integrity of the nigrostriatal system in Parkinson’s disease. Brain 136, 2419–2431.

4. Manson A, Stirpe P, and Schrag A (2012) Levodopa-induced-dyskinesias clinical features, incidence, risk factors, management and impact on quality of life. J Parkinsons Dis 2, 189–198.

5. Hechtner MC, Vogt T, Zollner Y, Schroder S, Sauer JB, Binder H, Singer S, and Mikolajczyk R (2014) Quality of life in Parkinson’s disease patients with motor fluctuations and dyskinesias in five European countries. Parkinsonism Relat Disord 20, 969–974.

6. Cenci MA, Riggare S, Pahwa R, Eidelberg D, and Hauser RA (2020) Dyskinesia matters. Mov Disord 35, 392–396.

7. Buck K, Voehringer P, and Ferger B (2010) Site-specific action of L-3,4-dihydroxyphenylalanine in the striatum but not globus pallidus and substantia nigra pars reticulata evokes dyskinetic movements in chronic L-3,4-dihydroxyphenylalanine-treated 6-hydroxydopamine-lesioned rats. Neuroscience 166, 355–358.

8. Buck K, and Ferger B (2008) Intrastriatal inhibition of aromatic amino acid decarboxylase prevents l-DOPA-induced dyskinesia: a bilateral reverse in vivo microdialysis study in 6-hydroxydopamine lesioned rats. Neurobiol Dis 29, 210–220.

9. Schrag A, and Quinn N (2000) Dyskinesias and motor fluctuations in Parkinson’s disease. A community-based study. Brain 123 ( Pt 11), 2297–2305.

10. Olanow WC, Kieburtz K, Rascol O, Poewe W, Schapira AH, Emre M, Nissinen H, Leinonen M, Stocchi F, and Stalevo Reduction in Dyskinesia Evaluation in Parkinson’s Disease I (2013) Factors predictive of the development of Levodopa-induced dyskinesia and wearing-off in Parkinson’s disease. Mov Disord 28, 1064–1071.

11. Olanow CW, Calabresi P, and Obeso JA (2020) Continuous Dopaminergic Stimulation as a Treatment for Parkinson’s Disease: Current Status and Future Opportunities. Mov Disord 35, 1731–1744.

12. Grandas F, Galiano ML, and Tabernero C (1999) Risk factors for levodopa-induced dyskinesias in Parkinson’s disease. J Neurol 246, 1127–1133.

13. Cilia R, Akpalu A, Sarfo FS, Cham M, Amboni M, Cereda E, Fabbri M, Adjei P, Akassi J, Bonetti A, and Pezzoli G (2014) The modern pre-levodopa era of Parkinson’s disease: insights into motor complications from sub-Saharan Africa. Brain 137, 2731–2742.

14. Borgkvist A, Lieberman OJ, and Sulzer D (2018) Synaptic plasticity may underlie l-DOPA induced dyskinesia. Curr Opin Neurobiol 48, 71–78.

15. Pons R, Syrengelas D, Youroukos S, Orfanou I, Dinopoulos A, Cormand B, Ormazabal A, Garzia-Cazorla A, Serrano M, and Artuch R (2013) Levodopa-induced dyskinesias in tyrosine hydroxylase deficiency. Mov Disord 28, 1058–1063.

16. Yoo HS, Chung SJ, Chung SJ, Moon H, Oh JS, Kim JS, Hong JY, Ye BS, Sohn YH, and Lee PH (2018) Presynaptic dopamine depletion determines the timing of levodopa-induced dyskinesia onset in Parkinson’s disease. Eur J Nucl Med Mol Imaging 45, 423–431.

17. Djaldetti R, Ziv I, and Melamed E (2006) The mystery of motor asymmetry in Parkinson’s disease. Lancet Neurol 5, 796–802.

18. Cedarbaum JM, Gandy SE, and McDowell FH (1991) “Early” initiation of levodopa treatment does not promote the development of motor response fluctuations, dyskinesias, or dementia in Parkinson’s disease. Neurology 41, 622–629.

19. Verschuur CVM, Suwijn SR, Boel JA, Post B, Bloem BR, van Hilten JJ, van Laar T, Tissingh G, Munts AG, Deuschl G, et al. (2019) Randomized Delayed-Start Trial of Levodopa in Parkinson’s Disease. N Engl J Med 380, 315–324.

20. Frequin HL, Verschuur CVM, Suwijn SR, Boel JA, Post B, Bloem BR, van Hilten JJ, van Laar T, Tissingh G, Munts AG, et al. (2024) Long-Term Follow-Up of the LEAP Study: Early Versus Delayed Levodopa in Early Parkinson’s Disease. Mov Disord 39, 975–982.

21. Ballard PA, Tetrud JW, and Langston JW (1985) Permanent human parkinsonism due to 1-methyl-4-phenyl-1,2,3,6-tetrahydropyridine (MPTP): seven cases. Neurology 35, 949–956.

22. Langston JW, and Ballard P (1984) Parkinsonism induced by 1-methyl-4-phenyl-1,2,3,6-tetrahydropyridine (MPTP): implications for treatment and the pathogenesis of Parkinson’s disease. Can J Neurol Sci 11, 160–165.

23. Langston JW, Ballard P, Tetrud JW, and Irwin I (1983) Chronic Parkinsonism in humans due to a product of meperidine-analog synthesis. Science 219, 979–980.

24. Shan L, Diaz O, Zhang Y, Ladenheim B, Cadet JL, Chiang YH, Olson L, Hoffer BJ, and Backman CM (2015) L-Dopa induced dyskinesias in Parkinsonian mice: Disease severity or L-Dopa history. Brain Res 1618, 261–269.

25. Winkler C, Kirik D, Bjorklund A, and Cenci MA (2002) L-DOPA-induced dyskinesia in the intrastriatal 6-hydroxydopamine model of parkinson’s disease: relation to motor and cellular parameters of nigrostriatal function. Neurobiol Dis 10, 165–186.

26. Servillo F, De Carluccio M, Di Lazzaro G, Campanelli F, Marino G, Natale G, Ledonne A, Massaro Cenere M, Paldino E, Di Giuda D, et al. (2024) Molecular and cellular determinants of L-Dopa-induced dyskinesia in Parkinson’s Disease. NPJ Parkinsons Dis 10, 228.

27. Pavon N, Martin AB, Mendialdua A, and Moratalla R (2006) ERK phosphorylation and FosB expression are associated with L-DOPA-induced dyskinesia in hemiparkinsonian mice. Biol Psychiatry 59, 64–74.

28. Andersson M, Hilbertson A, and Cenci MA (1999) Striatal fosB expression is causally linked with l-DOPA-induced abnormal involuntary movements and the associated upregulation of striatal prodynorphin mRNA in a rat model of Parkinson’s disease. Neurobiol Dis 6, 461–474.

29. Choi SJ, Ma TC, Ding Y, Cheung T, Joshi N, Sulzer D, Mosharov EV, and Kang UJ (2020) Alterations in the intrinsic properties of striatal cholinergic interneurons after dopamine lesion and chronic L-DOPA. Elife 9.

30. Cenci MA (2014) Presynaptic Mechanisms of l-DOPA-Induced Dyskinesia: The Findings, the Debate, and the Therapeutic Implications. Front Neurol 5, 242.

31. Giros B, Jaber M, Jones SR, Wightman RM, and Caron MG (1996) Hyperlocomotion and indifference to cocaine and amphetamine in mice lacking the dopamine transporter. Nature 379, 606–612.

32. Abercrombie ED, Bonatz AE, and Zigmond MJ (1990) Effects of L-dopa on extracellular dopamine in striatum of normal and 6-hydroxydopamine-treated rats. Brain Res 525, 36–44.

33. Rascol O, Nutt JG, Blin O, Goetz CG, Trugman JM, Soubrouillard C, Carter JH, Currie LJ, Fabre N, Thalamas C, et al. (2001) Induction by dopamine D1 receptor agonist ABT-431 of dyskinesia similar to levodopa in patients with Parkinson disease. Arch Neurol 58, 249–254.

34. Kim DS, Szczypka MS, and Palmiter RD (2000) Dopamine-deficient mice are hypersensitive to dopamine receptor agonists. J Neurosci 20, 4405–4413.

35. Gerfen CR (2003) D1 dopamine receptor supersensitivity in the dopamine-depleted striatum animal model of Parkinson’s disease. Neuroscientist 9, 455–462.

36. Kreitzer AC (2009) Physiology and pharmacology of striatal neurons. Annu Rev Neurosci 32, 127–147.

37. Tritsch NX, and Sabatini BL (2012) Dopaminergic modulation of synaptic transmission in cortex and striatum. Neuron 76, 33–50.

38. Bastide MF, Meissner WG, Picconi B, Fasano S, Fernagut PO, Feyder M, Francardo V, Alcacer C, Ding Y, Brambilla R, et al. (2015) Pathophysiology of L-dopa-induced motor and non-motor complications in Parkinson’s disease. Prog Neurobiol 132, 96–168.

39. McGregor MM, and Nelson AB (2019) Circuit Mechanisms of Parkinson’s Disease. Neuron 101, 1042–1056.

40. Espay AJ, Morgante F, Merola A, Fasano A, Marsili L, Fox SH, Bezard E, Picconi B, Calabresi P, and Lang AE (2018) Levodopa-induced dyskinesia in Parkinson disease: Current and evolving concepts. Ann Neurol 84, 797–811.

41. Picconi B, Hernandez LF, Obeso JA, and Calabresi P (2018) Motor complications in Parkinson’s disease: Striatal molecular and electrophysiological mechanisms of dyskinesias. Mov Disord 33, 867–876.

42. Cenci MA, and Kumar A (2024) Cells, pathways, and models in dyskinesia research. Curr Opin Neurobiol 84, 102833.

43. Kravitz AV, Freeze BS, Parker PR, Kay K, Thwin MT, Deisseroth K, and Kreitzer AC (2010) Regulation of parkinsonian motor behaviours by optogenetic control of basal ganglia circuitry. Nature 466, 622–626.

44. Gittis AH, Leventhal DK, Fensterheim BA, Pettibone JR, Berke JD, and Kreitzer AC (2011) Selective inhibition of striatal fast-spiking interneurons causes dyskinesias. J Neurosci 31, 15727–15731.

45. Kharkwal G, Brami-Cherrier K, Lizardi-Ortiz JE, Nelson AB, Ramos M, Del Barrio D, Sulzer D, Kreitzer AC, and Borrelli E (2016) Parkinsonism Driven by Antipsychotics Originates from Dopaminergic Control of Striatal Cholinergic Interneurons. Neuron 91, 67–78.

46. Sagot B, Li L, and Zhou FM (2018) Hyperactive Response of Direct Pathway Striatal Projection Neurons to L-dopa and D1 Agonism in Freely Moving Parkinsonian Mice. Front Neural Circuits 12, 57.

47. Paz R, Ryan M, Marcott P, Girasole A, Faryean J, Duong V, Sridhar S, and Nelson AB (2025) Repetitive Levodopa Treatment Drives Cell Type-Specific Striatal Adaptations Associated With Progressive Dyskinesia in Parkinsonian Mice. BioRxiv.

48. Maltese M, March JR, Bashaw AG, and Tritsch NX (2021) Dopamine differentially modulates the size of projection neuron ensembles in the intact and dopamine-depleted striatum. Elife 10.

49. Parker JG, Marshall JD, Ahanonu B, Wu YW, Kim TH, Grewe BF, Zhang Y, Li JZ, Ding JB, Ehlers MD, and Schnitzer MJ (2018) Diametric neural ensemble dynamics in parkinsonian and dyskinetic states. Nature 557, 177–182.

50. Ryan MB, Bair-Marshall C, and Nelson AB (2018) Aberrant Striatal Activity in Parkinsonism and Levodopa-Induced Dyskinesia. Cell Rep 23, 3438–3446 e3435.

51. Girasole AE, Lum MY, Nathaniel D, Bair-Marshall CJ, Guenthner CJ, Luo L, Kreitzer AC, and Nelson AB (2018) A Subpopulation of Striatal Neurons Mediates Levodopa-Induced Dyskinesia. Neuron 97, 787–795 e786.

52. Alcacer C, Andreoli L, Sebastianutto I, Jakobsson J, Fieblinger T, and Cenci MA (2017) Chemogenetic stimulation of striatal projection neurons modulates responses to Parkinson’s disease therapy. J Clin Invest 127, 720–734.

53. Castela I, Casado-Polanco R, Rubio YV, da Silva JA, Marquez R, Pro B, Moratalla R, Redgrave P, Costa RM, Obeso J, and Hernandez LF (2023) Selective activation of striatal indirect pathway suppresses levodopa induced-dyskinesias. Neurobiol Dis 176, 105930.

54. Keifman E, Ruiz-DeDiego I, Pafundo DE, Paz RM, Solis O, Murer MG, and Moratalla R (2019) Optostimulation of striatonigral terminals in substantia nigra induces dyskinesia that increases after L-DOPA in a mouse model of Parkinson’s disease. Br J Pharmacol 176, 2146–2161.

55. Rothwell PE, Hayton SJ, Sun GL, Fuccillo MV, Lim BK, and Malenka RC (2015) Input-and Output-Specific Regulation of Serial Order Performance by Corticostriatal Circuits. Neuron 88, 345–356.

56. Darmopil S, Martin AB, De Diego IR, Ares S, and Moratalla R (2009) Genetic inactivation of dopamine D1 but not D2 receptors inhibits L-DOPA-induced dyskinesia and histone activation. Biol Psychiatry 66, 603–613.

57. Westin JE, Vercammen L, Strome EM, Konradi C, and Cenci MA (2007) Spatiotemporal pattern of striatal ERK1/2 phosphorylation in a rat model of L-DOPA-induced dyskinesia and the role of dopamine D1 receptors. Biol Psychiatry 62, 800–810.

58. Mela F, Marti M, Bido S, Cenci MA, and Morari M (2012) In vivo evidence for a differential contribution of striatal and nigral D1 and D2 receptors to L-DOPA induced dyskinesia and the accompanying surge of nigral amino acid levels. Neurobiol Dis 45, 573–582.

59. Dobbs LK, Kaplan AR, Lemos JC, Matsui A, Rubinstein M, and Alvarez VA (2016) Dopamine Regulation of Lateral Inhibition between Striatal Neurons Gates the Stimulant Actions of Cocaine. Neuron 90, 1100–1113.

60. Yun S, Yang B, Anair JD, Martin MM, Fleps SW, Pamukcu A, Yeh NH, Contractor A, Kennedy A, and Parker JG (2023) Antipsychotic drug efficacy correlates with the modulation of D1 rather than D2 receptor-expressing striatal projection neurons. Nat Neurosci 26, 1417–1428.

61. Isett BR, Nguyen KP, Schwenk JC, Yurek JR, Snyder CN, Vounatsos MV, Adegbesan KA, Ziausyte U, and Gittis AH (2023) The indirect pathway of the basal ganglia promotes transient punishment but not motor suppression. Neuron 111, 2218–2231 e2214.

62. Gertler TS, Chan CS, and Surmeier DJ (2008) Dichotomous anatomical properties of adult striatal medium spiny neurons. J Neurosci 28, 10814–10824.

63. West AE, Griffith EC, and Greenberg ME (2002) Regulation of transcription factors by neuronal activity. Nat Rev Neurosci 3, 921–931.

64. Pettit NL, Yap EL, Greenberg ME, and Harvey CD (2022) Fos ensembles encode and shape stable spatial maps in the hippocampus. Nature 609, 327–334.

65. Cruz-Mendoza F, Jauregui-Huerta F, Aguilar-Delgadillo A, Garcia-Estrada J, and Luquin S (2022) Immediate Early Gene c-fos in the Brain: Focus on Glial Cells. Brain Sci 12.

66. Steward O, and Worley PF (2001) Selective targeting of newly synthesized Arc mRNA to active synapses requires NMDA receptor activation. Neuron 30, 227–240.

67. Bramham CR, Worley PF, Moore MJ, and Guzowski JF (2008) The immediate early gene arc/arg3.1: regulation, mechanisms, and function. J Neurosci 28, 11760–11767.

68. Azdad K, Chavez M, Don Bischop P, Wetzelaer P, Marescau B, De Deyn PP, Gall D, and Schiffmann SN (2009) Homeostatic plasticity of striatal neurons intrinsic excitability following dopamine depletion. PLoS One 4, e6908.

69. Maurice N, Liberge M, Jaouen F, Ztaou S, Hanini M, Camon J, Deisseroth K, Amalric M, Kerkerian-Le Goff L, and Beurrier C (2015) Striatal Cholinergic Interneurons Control Motor Behavior and Basal Ganglia Function in Experimental Parkinsonism. Cell Rep 13, 657–666.

70. Fieblinger T, Graves SM, Sebel LE, Alcacer C, Plotkin JL, Gertler TS, Chan CS, Heiman M, Greengard P, Cenci MA, and Surmeier DJ (2014) Cell type-specific plasticity of striatal projection neurons in parkinsonism and L-DOPA-induced dyskinesia. Nat Commun 5, 5316.

71. Ketzef M, Spigolon G, Johansson Y, Bonito-Oliva A, Fisone G, and Silberberg G (2017) Dopamine Depletion Impairs Bilateral Sensory Processing in the Striatum in a Pathway-Dependent Manner. Neuron 94, 855–865 e855.

72. Melani R, and Tritsch NX (2025). Beyond dopamine: Cotransmission by dopamine neurons. In The Handbook of Dopamine, S.J. Cragg, and M. Walton, eds. (Elsevier), pp. 141–151.

73. Melani R, and Tritsch NX (2022) Inhibitory co-transmission from midbrain dopamine neurons relies on presynaptic GABA uptake. Cell Rep 39, 110716.

74. Trudeau LE, Hnasko TS, Wallen-Mackenzie A, Morales M, Rayport S, and Sulzer D (2014) The multilingual nature of dopamine neurons. Prog Brain Res 211, 141–164.

75. Malave L, Zuelke DR, Uribe-Cano S, Starikov L, Rebholz H, Friedman E, Qin C, Li Q, Bezard E, and Kottmann AH (2021) Dopaminergic co-transmission with sonic hedgehog inhibits abnormal involuntary movements in models of Parkinson’s disease and L-Dopa induced dyskinesia. Commun Biol 4, 1071.

76. Singh A (2018) Oscillatory activity in the cortico-basal ganglia-thalamic neural circuits in Parkinson’s disease. Eur J Neurosci 48, 2869–2878.

77. Willard AM, Isett BR, Whalen TC, Mastro KJ, Ki CS, Mao X, and Gittis AH (2019) State transitions in the substantia nigra reticulata predict the onset of motor deficits in models of progressive dopamine depletion in mice. Elife 8.

78. Sarinana J, Kitamura T, Kunzler P, Sultzman L, and Tonegawa S (2014) Differential roles of the dopamine 1-class receptors, D1R and D5R, in hippocampal dependent memory. Proc Natl Acad Sci U S A 111, 8245–8250.

79. Farrell MS, Pei Y, Wan Y, Yadav PN, Daigle TL, Urban DJ, Lee HM, Sciaky N, Simmons A, Nonneman RJ, et al. (2013) A Galphas DREADD mouse for selective modulation of cAMP production in striatopallidal neurons. Neuropsychopharmacology 38, 854–862.

80. Nagai Y, Miyakawa N, Takuwa H, Hori Y, Oyama K, Ji B, Takahashi M, Huang XP, Slocum ST, DiBerto JF, et al. (2020) Deschloroclozapine, a potent and selective chemogenetic actuator enables rapid neuronal and behavioral modulations in mice and monkeys. Nat Neurosci 23, 1157–1167.

81. Curtze C, Nutt JG, Carlson-Kuhta P, Mancini M, and Horak FB (2015) Levodopa Is a Double-Edged Sword for Balance and Gait in People With Parkinson’s Disease. Mov Disord 30, 1361–1370.

82. Kish SJ, Shannak K, and Hornykiewicz O (1988) Uneven pattern of dopamine loss in the striatum of patients with idiopathic Parkinson’s disease. Pathophysiologic and clinical implications. N Engl J Med 318, 876–880.

83. Damier P, Hirsch EC, Agid Y, and Graybiel AM (1999) The substantia nigra of the human brain. II. Patterns of loss of dopamine-containing neurons in Parkinson’s disease. Brain 122 ( Pt 8), 1437–1448.

84. Gerfen CR, Miyachi S, Paletzki R, and Brown P (2002) D1 dopamine receptor supersensitivity in the dopamine-depleted striatum results from a switch in the regulation of ERK1/2/MAP kinase. J Neurosci 22, 5042–5054.

85. Druart M, Yang YC, Tritsch NX, and Sippy T (2026) The Cellular and Synaptic Actions of Dopamine During Behavior. BioRxiv.

86. Saunders A, Macosko EZ, Wysoker A, Goldman M, Krienen FM, de Rivera H, Bien E, Baum M, Bortolin L, Wang S, et al. (2018) Molecular Diversity and Specializations among the Cells of the Adult Mouse Brain. Cell 174, 1015–1030 e1016.

87. Francardo V, Recchia A, Popovic N, Andersson D, Nissbrandt H, and Cenci MA (2011) Impact of the lesion procedure on the profiles of motor impairment and molecular responsiveness to L-DOPA in the 6-hydroxydopamine mouse model of Parkinson’s disease. Neurobiol Dis 42, 327–340.

88. Soileau MJ, Aldred J, Budur K, Fisseha N, Fung VS, Jeong A, Kimber TE, Klos K, Litvan I, O’Neill D, et al. (2022) Safety and efficacy of continuous subcutaneous foslevodopa-foscarbidopa in patients with advanced Parkinson’s disease: a randomised, double-blind, active-controlled, phase 3 trial. Lancet Neurol 21, 1099–1109.

89. Freire-Alvarez E, Kurca E, Lopez Manzanares L, Pekkonen E, Spanaki C, Vanni P, Liu Y, Sanchez-Solino O, and Barbato LM (2021) Levodopa-Carbidopa Intestinal Gel Reduces Dyskinesia in Parkinson’s Disease in a Randomized Trial. Mov Disord 36, 2615–2623.

90. Olanow CW, Kieburtz K, Odin P, Espay AJ, Standaert DG, Fernandez HH, Vanagunas A, Othman AA, Widnell KL, Robieson WZ, et al. (2014) Continuous intrajejunal infusion of levodopa-carbidopa intestinal gel for patients with advanced Parkinson’s disease: a randomised, controlled, double-blind, double-dummy study. Lancet Neurol 13, 141–149.

91. Kim DS, Palmiter RD, Cummins A, and Gerfen CR (2006) Reversal of supersensitive striatal dopamine D1 receptor signaling and extracellular signal-regulated kinase activity in dopamine-deficient mice. Neuroscience 137, 1381–1388.

92. Liu H, Melani R, Maltese M, Taniguchi J, Sankaramanchi A, Zeng R, Martin JR, and Tritsch NX (2025) Subsecond dopamine fluctuations do not specify the vigor of ongoing actions. Nature Neuroscience.

